# Multiplexed single-cell profiling of chromatin states at genomic loci by expansion microscopy

**DOI:** 10.1101/2020.11.17.385476

**Authors:** Marcus A. Woodworth, Kenneth K.H. Ng, Aaron R. Halpern, Nicholas A. Pease, Phuc H.B. Nguyen, Hao Yuan Kueh, Joshua C. Vaughan

**Author notes:** Co-corresponding authors: H.Y.K., J.C.V.

## Abstract

Proper regulation of genome architecture and activity is essential for the development and function of multicellular organisms. Histone modifications, acting in combination, specify these activity states at individual genomic loci. However, the methods used to study these modifications often require either a large number of cells or are limited to targeting one histone mark at a time. Here, we developed a new method called Single Cell Evaluation of Post-TRanslational Epigenetic Encoding (SCEPTRE) that uses Expansion Microscopy (ExM) to visualize and quantify multiple histone modifications at non-repetitive genomic regions in single cells at a spatial resolution of ~75 nm. Using SCEPTRE, we distinguished multiple histone modifications at a single housekeeping gene, quantified histone modification levels at multiple developmentally-regulated genes in individual cells, and identified a relationship between histone H3K4 trimethylation and the loading of paused RNA polymerase II at individual loci. Thus, SCEPTRE enables multiplexed detection of combinatorial chromatin states at single genomic loci in single cells.

## Introduction

Proper regulation of genome activity and architecture is critical for development, growth, and function of a multicellular organism.^1,2^ Regulation occurs in large part at the nucleosome, where ~147 base pairs of DNA wrap around an octamer of 4 different histone pairs: H2A, H2B, H3 and H4.^3^ Various residues found at the N and C-terminal tails of these histones can acquire post-translational modifications, such as acetylation and methylation, which grant nucleosomes the ability to either participate in organized compaction of chromatin or to recruit transcriptionally relevant protein complexes.^4,5^ Researchers have therefore suggested that these modifications, also known as histone marks, act as a code for the epigenetic state of genomic regions.^6,7^ Although several sequencing-based methods are available for studying distinct histone modifications (i.e., ChIP-seq),^8,9^ chromatin accessibility,^10,11^ genomic contact frequencies,^12,13^ and genomic nuclear locations,^14^ these methods are either unable to resolve cell-to-cell variations or are limited to studying one histone modification at a time. Therefore, the role these marks play in controlling chromatin structure and gene expression at the single cell and single locus level remains poorly understood and vigorously debated.

To tackle this problem, super-resolution fluorescence microscopy techniques have been used to observe more closely how histone marks impact chromatin organization within a cell’s nucleus. Using Stochastic Optical Reconstruction Microscopy (STORM),^15,16^ researchers saw that nucleosomes form clusters that vary in size and nuclear distribution depending on a cell’s developmental stage or what histone marks they present.^17,18^ Others have combined STORM with DNA Fluorescence *in situ* hybridization (FISH) to map spatial aspects of genomic loci with a spatial resolution comparable to the observed sizes of these histone clusters.^19^ Collectively, these studies suggest that concurrent visualization of DNA and histone modifications with super-resolution microscopy could enable profiling chromatin states at the level of single loci. However, most studies to date have viewed histone marks and genes separately, because combining immunofluorescence and DNA FISH can be challenging due to the harsh solvents and/or high temperatures used in FISH protocols.^20–23^ Although researchers have visualized immunolabeled histone marks across whole chromosomes,^21,22^ or at repetitive and highly abundant ALU elements regions labeled with an alternative hybridization strategy,^24^ there are still no methods available to study multiple histone marks at individual non-repetitive genomic loci within a nucleus. A better understanding of histone mark heterogeneity at individual loci would require a new method capable of further decoupling immunofluorescence and FISH labeling.

We therefore developed a new method, called Single Cell Evaluation of Post-TRanslational Epigenetic Encoding (SCEPTRE), which uses expansion microscopy (ExM)^25,26^ to combine DNA FISH with immunofluorescence and quantify histone mark fluorescence signals at individual loci within the nucleus. ExM preserves the signal of antibody labels on protein structures by covalently linking antibodies and proteins to a swellable hydrogel that is grown within the sample.^25,26^ This signal preservation enables subsequent use of relatively harsh conditions, such as high temperatures and organic solvents, for labeling of genomic DNA by FISH without loss of the antibody signal. At the same time, ExM enables the isotropic expansion of specimens with low distortion so that these specimens may be examined with a high spatial resolution (here ~75 nm) in the expanded state even when using conventional microscopes with a diffraction-limited resolution of ~250 nm. We demonstrate the capabilities of SCEPTRE for a variety of systems: 1) we compared signals of multiple histone marks at a housekeeping gene locus; 2) We distinguished histone mark signals between developmentally-regulated genes in a single cell; 3) we demonstrate a correlation between histone marks and paused RNA polymerase II in a single region. Together, these experiments establish SCEPTRE as a powerful tool to study the role histone marks have at individual genes within the nuclei of single cells.

## Materials and Methods

### Reagents

The following primary antibodies were purchased and used for immunofluorescence: Human anti-centromeres (Antibodies Incorporated, 15-235), Mouse anti-H3K27me3 (Active Motif, 61017), Mouse anti-H3K4me3 (EMD Millipore, 05-1339-S), Mouse anti-RNA polymerase II CTD repeat YSPTSPS phosphorylated at Serine 5 (Abcam, ab5408). The following primary antibodies were purchased and used for immunofluorescence and CUT&RUN^27,28^ followed by sequencing: Rabbit anti-H3K4me3 (Active motif, 39159), Rabbit anti-H3K27me3 (Active Motif, 39155), Rabbit anti-H3K27ac (Active Motif, 39133). The following unconjugated secondary antibodies were purchased from Jackson ImmunoResearch: Donkey anti-rabbit (711-005-152) and Donkey anti-human (709-005-149). The following conjugated secondary antibodies were purchased from Jackson ImmunoResearch: Donkey anti-rabbit conjugated with Alexa Fluor 488 (711-545-152) and Donkey anti-mouse conjugated with Alexa Fluor 488 (715-545-150).

The following enzymes were purchased: proteinase K (Thermo Fisher Scientific, EO0491), RNase A (Thermo Fisher Scientific, EN0531), alcohol oxidase (Sigma-Aldrich, A2404-1KU), catalase (Sigma-Aldrich, C100), Phusion Hot-start master mix (New England Biolabs, M0536L), DNase I (New England Biolabs, M0303A) and Maxima H Minus RT Transcriptase (Thermo Fisher Scientific, EP0752).

The following chemical reagents were purchased: 10× phosphate-buffered saline (PBS, Fisher Bioreagents, BP399-1), 32% paraformaldehyde aqueous solution (PFA, Electron Microscopy Sciences, RT15714), 4-(1,1,3,3-tetramethylbutyl)phenyl-polyethylene glycol (Triton X-100, Sigma-Aldrich, X100), Bovine serum albumin (BSA, Rockland Immunochemicals Inc., BSA-50), ATTO 488 NHS-ester (ATTO-TEC GmbH, AD 488-35), Alexa Fluor 568 NHS-ester (Thermo Fisher Scientific, A-20003), methacrylic acid NHS-ester (MA-NHS, Sigma-Aldrich, 730300), 40% acrylamide aqueous solution (Bio-Rad Laboratories, 1610140), 2% bis-acrylamide aqueous solution (Bio-Rad Laboratories, 1610142), 97% sodium acrylate powder (Sigma-Aldrich, 408220), ammonium persulfate (APS, Thermo Fisher Scientific, 17874), tetramethylethylenediamine (TEMED, Thermo Fisher Scientific, 17919), 10× tris-acetate-EDTA (TAE, Fisher Bioreagents, BP2434-4), guanidine hydrochloride powder (Sigma-Aldrich, G3272), sodium azide (Sigma-Aldrich, S2002), poly-L-lysine (Sigma-Aldrich, P8920), sodium bicarbonate (VWR, 470302), formamide (Fisher Chemical, F84-1), 20× saline sodium citrate (SSC, Sigma-Aldrich, S6639), 50% OmniPur Dextran Sulfate (EMD Millipore, 3730), Tween 20 (Sigma-Aldrich, P9416), Hoechst 33258 (Sigma-Aldrich, B2883-25MG), Tris Base (Fisher scientific, BP152-500), methyl viologen dichloride hydrate (Sigma-Aldrich 856177), L-ascorbic acid (Fisher scientific, A61-25), digitonin (EMD Millipore, 300410), glycogen (VWR, 97063-256), sodium chloride (NaCl, Thermo Fisher Scientific, S271500), Ethylenediaminetetraacetic acid disodium salt dihydrate (EDTA, Sigma-Aldrich, E6635), Ethylene glycol-bis(2-aminoethylether)-N,N,N’,N’-tetraacetic acid (EGTA, Sigma-Aldrich, E4378) and calcium chloride dihydrate (VWR, 0556).

Alpha-satellite, GAPDH set, adapter and conjugated reporter oligonucleotide probes were obtained from Integrated DNA Technologies (IDT). A Precise Synthetic Oligo Pool (SC1966-12) containing probes covering the *MYL6, HOXC* and *LINC-PINT* regions was obtained from GenScript (for a list of sequences, see supplementary spreadsheet).

### Cell culture

h-TERT RPE1 cells were cultured and grown to ~80% confluency using Dulbecco’s modified eagle medium (Gibco, 11995065) supplemented with 100 units/mL of penicillin and streptomycin (Gibco, 15140122), 1% nonessential amino acids (Gibco, 11140050), and 10% fetal bovine serum (Gibco, 26140079). Cells were then trypsinized with 0.25% trypsin-EDTA (Gibco, 25200056) and seeded at ~75,000 cells per well on top of round coverslips (no. 1.5, ~12 mm diameter) placed within 24-well culture plates. After growing overnight (~18 hours), the cells were briefly rinsed with 1× PBS then fixed with either 4-10% PFA in 1× PBS for 10 minutes at room temperature (~22 °C), or in cold EtOH:MeOH (1:1) for 6 minutes at - 20 °C. Fixed cells were washed three times with 1× PBS, then stored in 1× PBS azide (1 × PBS with 3 mM sodium azide) at 4 °C before use (see **sup. table 1** for more details).

### Secondary antibody fluorophore conjugation

Conjugation was performed by mixing 40 μL of a secondary antibody solution with 5 μL of a 1 M sodium bicarbonate solution, then adding 2-5 μg of an NHS ester functionalized fluorophore. The mixture was left to react for 30 minutes protected from ambient light and the crude reaction mixture was passed through a NAP-5 column (GE Healthcare Life Sciences, 17085301) for collection and purification of the fluorophore-conjugated secondary antibody. Further characterization of the secondary antibody was done by ultraviolet/visible absorption spectroscopy.

### Immunostaining procedure

The immunostain procedure was adapted from previous protocols,^17,18^ and goes as follows: fixed RPE1 cells were incubated first in permeabilization solution (1× PBS with 0.1% (v/v) TritonX-100) for 10 minutes, then washed three times with 1× PBS. After permeabilization, cells were incubated in block solution (1× PBS with 10% (w/v) BSA and 3 mM sodium azide) for 1 hour at room temperature, followed by incubation in primary solution (2-5 μg/mL of primary antibodies diluted in block) overnight at 4 °C. The sample was washed with block three times (10 minutes each time), then incubated in secondary solution (2-3 μg/mL of secondary fluorophore-conjugated antibodies in block) for 1-2 hours at room temperature. The sample was washed once for 10 minutes with block, then three times with 1× PBS azide. Samples which had been originally fixed in EtOH:MeOH were post-fixed in 4% PFA in 1× PBS for 10 minutes, then washed three times with 1× PBS azide. Immunostained samples were either immediately gelled or stored in 1× PBS azide at 4 °C for up to ~1 week for later use (see **sup. table 1** for more details).

### Cell gelation, digestion, and expansion

Expansion microscopy was adapted from a previous protocol,^26^ and goes as follows: Immunolabeled cells were treated with freshly prepared 5 mM MA-NHS in 1× PBS for 10 minutes, then washed three times with 1× PBS. Cells were incubated in monomer solution (1× PBS with 2 M NaCl, 2.5% (w/w) acrylamide, 0.15% (w/w) N,N’-methylenebisacrylamide and 8.625% (w/w) sodium acrylate) for 10 minutes before gelation with 0.15-0.2% (w/v) APS and 0.2% TEMED (w/w) at room temperature for at least 30 minutes in a sealed container backfilled with nitrogen gas. After polymerization, the cell-embedded hydrogel was gently removed from the 12 mm coverslip, then incubated in digestion solution (1× TAE with 0.5% (v/v) Triton X-100, 0.8 M guanidine HCl and 8 units/mL proteinase K) overnight at 37 °C. The digested sample was both washed and expanded by placing the sample in deionized water, which was replaced every 15-20 minutes for at least three times. Hydrogels were stored in 2× SSC at 4 °C, typically up to ~1 month.

### DNA fluorescence in situ hybridization

The general DNA FISH procedure for non-repetitive genomic regions (*GAPDH, MYL6, HOXC* and *LINC-PINT*) was adapted from previous protocols,^29,30^ and goes as follows: Briefly, a small (~20 μL) piece of gel from each expanded cell sample was first incubated in hybridization buffer (2× SSC with 50% (v/v) formamide and 0.1% (v/v) Tween 20) for 10 minutes at room temperature. Samples were incubated in pre-heated hybridization buffer for 30 minutes at 60 °C. A hybridization mixture (2× SSC with 50% formamide (v/v), 10% dextran sulfate (w/v), 0.1% (v/v) Tween 20, 3 mM sodium azide, ~10-20 nM oligo probe library per kb of targeted genomic region, and 1-1.5× concentration of oligo reporters and adapters to oligo probe library) specific to each sample was preheated to 90 °C for 5-10 minutes and then added to each sample at an approximate 2:1 volume ratio. Samples were denatured at 90-92.5 °C for 2.5-10 minutes and hybridized at 37-42 °C overnight. Samples were washed three times, 15 minutes each time: first with preheated 2× SSCT (2× SSC with 0.1% (v/v) Tween 20) at 60 °C, then with preheated 2× SSCT at 37 °C, and lastly with 2× SSCT at room temperature. Samples were stored at 4 °C in 0.2× SSCT (0.2× SSC with 0.01% (v/v) Tween 20) until needed (within a week). Samples were fully expanded to ~4× the original size with deionized water at 4 °C, replacing the water twice every 10 minutes (see **sup. table 1** for more details).

The DNA FISH procedure for the repetitive alpha-satellite region was done as follows: Expanded RPE1 cells were incubated for 1 hour at room temperature in 1× PBS. The sample was then incubated in 1× PBS supplemented with 100 μg/mL of Rnase A for 1 hour at 37 °C. After RNA digestion, the sample was incubated in 2× SSCT for 30 minutes at room temperature. The samples were then incubated in hybridization buffer for 30 minutes at room temperature. The gel was transferred to a hybridization buffer containing 200nM of alpha-satellite oligonucleotide probe. The sample was denatured for 15 minutes at 95 °C. Gels were washed once in 20× SSC for 15 minutes at 37 °C, then in 2× SSC for 1 hour at 37 °C. The samples were incubated in 2× SSC with 200 nM alpha-satellite adapter probe and 600 nM of reporter probe A for 30 minutes at 37 °C. The sample was washed with 20× SSC for 20 minutes at 37 °C and lastly with 2× SSC for 20 minutes at room temperature. After this, the alpha-satellite sample was expanded to ~3× the original size by incubating the sample in 0.2× SSC, then a second time in 0.2× SSC with 1 μg/mL of Hoechst 33258 (see **sup. table 1** for more details).

### Sample mounting and imaging

For expanded samples using Alexa Fluor 750 fluorophore-conjugated reporters, samples were incubated in imaging buffer (10 mM Tris buffer (pH 8) with 1 mM Methyl viologen, 1 mM Ascorbic acid and 2% (v/v) MeOH) for 10 minutes. Before imaging, samples were first adhered to a poly-L-lysine–coated rectangular no. 1.5 coverslip, then they were supplemented with ~30 units/mL alcohol oxidase and 0.2% (w/v) catalase. Samples that did not have Alexa Fluor 750 were adhered to a poly-L-lysine–coated rectangular no. 1.5 coverslip. All samples were imaged either with a Leica SP5 inverted confocal point scanning microscope at the University of Washington Biology Imaging Facility with a Plan Apo CS 63×, 1.2 numerical aperture (NA) water-immersion objective, or with a homebuilt spinning disk confocal microscope using a Nikon CFI60 Plan Apochromat 60×, 1.27 NA (Nikon) water immersion lens.

### CUT&RUN H3K4me3, H3K27me3 and H3K27ac profiling

CUT&RUN was performed as previously described,^28^ with the following adaptations: 250,000 trypsinized RPE1 cells were used per antibody condition. Cells were bound to Concanavalin A coated magnetic beads (Bangs Laboratories, BP531), permeabilized with 0.025% (w/v) digitonin, then incubated overnight with 5 μg of either anti-H3K4me3 (Active Motif, 39159), anti-H3K27me3 (Active Motif, 39155) or anti-H3K27ac (Active Motif, 39133) at 4 °C. Cells were washed then incubated with protein A-MNase fusion protein (a gift from S. Henikoff, FHCRC) for 15 minutes at room temperature. After another wash, cells were incubated with 2 mM calcium chloride for 30 minutes at 0 °C to induce MNase cleavage activity. The reaction was stopped with 2× STOP buffer (200 mM NaCl, 20 mM EDTA, 4 mM EGTA, 50 μg/mL RNase A, 50 μg/mL glycogen) and 0.2 pg of yeast spike-in DNA was added to each sample. Cleaved Histone-DNA complexes were isolated by centrifugation and DNA was extracted with a NucleoSpin PCR Clean-up kit (Macherey-Nagel, 740609).

Library preparation for each CUT&RUN antibody condition was done with a KAPA Hyper Prep Kit (VWR, 89125-040) with the PCR amplification settings adjusted to have simultaneous annealing and extension steps at 60 °C for 10 seconds. Library products between 200-300 base pairs were selected using Agencourt AMPure XP beads (Beckman Coulter, A63880) then sequenced with an Illumina MiSeq system at the University of Washington Northwest Genomics Center with paired-end 25 base pair sequencing read length and TruSeq primer standard for ~6 million reads per condition.

Paired-end sequencing reads were aligned separately to human (GRCh38/hg38) and yeast genomes using Bowtie2^31^ with the previously suggested specifications for mapping CUT&RUN sequencing data:^28^ --local --very-sensitive-local --no-unal --no-mixed --no-discordant −I 10 −X 700. Alignment results were converted to BAM files with SAMtools^32^ and then to BED files with BEDTools.^33^ Reads were sorted and filtered to remove random chromosomes, then, with BEDTools genomecov, histograms were generated for the mapped reads using spiked-in yeast reads and the number of cells for each condition as scaling factors. The results were visualized using the WashU Epigenome Browser (https://epigenomegateway.wustl.edu/).^34^

### Oligonucleotide probe design and amplification

DNA FISH probes were designed using OligoMiner,^35^ with standard buffer, length and melting temperature conditions, with the exception of the target *MYL6,* which had the following adaptations: base length between 28-42 nucleotides and melting temperature between 38-46 °C. Unique DNA sequences, which were previously screened for DNA FISH purposes,^19,36^ were appended to each probe as adapter/reporter hybridizing regions specific to each gene, along with a primer set for amplification. Designed probes were purchased as part of an oligo pool from GenScript, and the probes were amplified using a T7/Reverse-Transcriptase amplification protocol previously published,^30^ in an RNase-free environment with the following adaptations: After PCR amplification with a Phusion Hot-start master mix and purification with a DNA Clean & Concentrator-5 kit (Zymo Research, D4013), probes were T7 amplified with a HiScribe T7 Quick High Yield RNA Synthesis Kit (New England BioLabs, E2050S) supplemented with 1.3 units/μL RNaseOUT (ThermoFisher Scientific, 10777019) for 16 hours at 37 °C. DNA was digested with DNase I for 1 hour at 37 °C. RNA was purified from the sample by first adding LiCl solution from the HiScribe Kit at a 1:7 ratio to the RNA solution, incubating the solution at - 20 °C for 30 minutes and pelleting the precipitated RNA by centrifugation (~17,000 g) for 15 minutes at 4 °C. The supernatant was removed from the tube and the pellet was washed with 70% EtOH. The RNA pellet was centrifuged (~17,000 g) for 5 minutes at 4 °C and, after carefully removing the supernatant, the pellet was left to dry for 3 minutes. The RNA was dissolved in water and ~50 μg of RNA was added to Maxima H Minus RT buffer with 2.86 units/μL Maxima H Minus RT Transcriptase, 2.3 units/μL RNaseOUT, 1 mM dNTP and 14 μM Forward project primer. The solution was incubated at 50 °C for 2 hours, then samples were digested with 100 μg/mL RNase A for 1 hour at 37 °C. After RNase digestion, oligonucleotide probes were purified using a DNA Clean & Concentrator-25 kit (Zymo Research, D4033) with Oligo binding buffer (Zymo Research, D4060-1-10). The final product was assumed to have full yield (for a list of sequences, see supplementary spreadsheet).

### Image processing and analysis for SCEPTRE profiling

Image processing and analysis was performed using MATLAB. First, raw images obtained from immunofluorescence channels were smoothed with a gaussian filter using 1-2 standard deviations within a 3×3×3 matrix. Smoothed images were contrast adjusted, where background pixel levels were clipped at an adaptively determined threshold for each image set at 2-9 third quartiles away from the median of each image stack histogram. The contrast adjusted images were binarized, either by an Otsu method or a Laplace filter with alpha=0.2 followed by selection of all negative values. A nuclear mask (generated as described below) was applied to the binarized immunofluorescence channel and, after small components (volume < 20 voxels) were removed, a watershed was applied to the segmented clusters. Features, including mean fluorescence intensity of every immunofluorescence channel and overlap with clusters of other segmented immunofluorescence channels, were identified for each segmented cluster (see **sup. table 2** for more details). For the first image stack, each step was visually inspected to confirm proper threshold levels.

The nuclear mask was generated by applying the same segmentation process from above to either a Hoechst stain channel or the same immunolabeled channel with a contrast adjustment done with 1-3 third quartiles clipping. The segmented channel was subject to morphological opening with a sphere of 3 pixel radius, to fuse clusters within the nucleus. A convex hull was applied to the largest component (i.e. the nucleus) after it was selected from the rest of clusters. The segmented nucleus was morphologically closed with the same sphere that was used to morphologically open the channel (see **sup. table 2** for more details). For the first image stack, each step was visually inspected to confirm proper threshold levels.

After segmentation of the nuclear channel and immunofluorescence channels, the FISH raw channel was segmented in the same manner with the following exceptions: 1) The nuclear mask was applied before smoothing and contrast adjustment; 2) Clipping during contrast adjustment was performed with a threshold of 10-15 third quartiles away from the median; 3) No watershed was applied to FISH segmented regions; 4) clusters intersecting the periphery of the nuclear mask (e.g. highly fluorescent contaminant in FISH channel next to nucleus) were removed. Features, including mean fluorescence intensity of every immunofluorescence channel and overlap with each immunofluorescence segmented cluster, were identified for each segmented FISH cluster. Since the segmented FISH channel can contain small and dim clusters that, by visual inspection, do not correspond to the FISH-labeled genomic loci, small clusters (volume < 20-80 voxels) were filtered out before analyses. After segmentation of the FISH channel, randomly selected cubic regions were generated throughout the nuclear region of each image stack with a volume approximately equal to the mean volume of the selected FISH clusters. Mean fluorescence intensities of each immunofluorescence channel were determined for these random clusters (see **sup. table 2** for more details). For the first image stack, each step was visually inspected to confirm proper threshold levels.

Data obtained from the segmented clusters were inspected using contour and scatter plots with MATLAB built-in functions, or violin plots, using the MATLAB script violinplot (https://github.com/bastibe/Violinplot-Matlab). Contours were smoothed with a gaussian filter using 1 standard deviation within a 5×5 matrix. Correlation coefficients were determined using the MATLAB function corrcoef.

### Statistical analyses

Each figure, along with their related supplementary figures, represents an individual experiment where all cells were labeled, expanded, imaged and processed under the same conditions. Cell numbers for each experiment were: 1 (**fig. 2**), 50 (**fig. 3, sup. fig. 5, 6**), 52 (**fig. 4A, sup. fig. 7A, 8A**), 38 (**fig. 4B, sup. fig. 7B, 8B**), 54 (**fig. 5, sup. fig. 10, 11**), 1 (**sup. fig. 1**), 36 (**sup. fig. 3A**), 10 (**sup. fig. 3B**), 48 (**sup. fig. 4**), 40 (**sup. fig. 12–13**), 20 (**sup. fig. 14–15**).

Fluorescence signal, defined as the mean fluorescence intensity for a given immunolabeled channel within a cluster of the same experiment, was used as the main measurement for comparing histone mark or paused RNA polymerase II levels within the segmented clusters of immunolabeled, FISH-labeled or randomly selected regions, or between the distribution of fluorescence signals for each set of clusters. An arbitrary “on” threshold (equal to the 5th percentile of the fluorescent signal found within a respective immunolabeled cluster set) is represented in all graphs excluding violin plots, as a qualitative determinant of high or low fluorescence signal within each set of clusters. Correlation coefficients were determined for each comparison between fluorescence signals within a set of clusters. A right-tailed Wilcoxon rank sum test was used to determine if for a given experiment the median fluorescence signal of a cluster set was significantly higher than the median signal in randomly selected regions or a separate set. All numbers corresponding to fraction of overlap and distance are represented as mean ± standard deviation.

### Data availability

CUT&RUN sequencing data for H3K4me3, H3K27me3 and H3K27ac was submitted to the NCBI gene expression omnibus (http://www.ncbi.nlm.nih.gov/geo/) under the accession number GSE160784. Additional data related to this paper will be made available by the corresponding authors upon reasonable request.

## Results

### SCEPTRE uses ExM to co-localize immunolabeled proteins at DNA FISH labeled genomic regions

The labeling of individual genomic loci by DNA fluorescence *in situ* hybridization (FISH) has provided a powerful tool for visualizing chromatin structure in single cells.^29,37–39^ While DNA FISH could be combined with immunofluorescence labeling to enable concurrent visualization of chromatin modification states and associated proteins, integration of these two techniques has been challenging, because the harsh conditions required to melt double-stranded genomic DNA during labeling (e.g., treatment with hot formamide) may remove antibody labels applied before FISH or may compromise the antigenicity of relevant epitopes for post-FISH immunolabeling.^20–23^ To overcome this challenge, we employed expansion microscopy (ExM) as a means to preserve the signal of immunolabeled protein structures during DNA FISH labeling. In ExM, immunolabeled structures are covalently linked to a swellable hydrogel polymer scaffold that is isotropically expanded in deionized water in order to reveal features closer than the ~250 nm diffraction limit of light in the expanded state.^25,26^ ExM not only provides a high spatial resolution (~75 nm or better when using a standard confocal microscope with ~4× expanding gels), but also enables antibody labels to be covalently tethered to the hydrogel scaffold, such that DNA FISH can subsequently be performed without loss of antibody fluorescence. ExM has previously been combined with DNA FISH to either visualize the *HER2* gene in tissue,^40^ or to visualize repetitive centromere regions in plants.^41^ However this combination has not yet been used to determine the density of a protein structure, such as histone mark clusters, at specific genomic regions. We refer to this new methodology as Single Cell Evaluation of Post-TRanslational Epigenetic Encoding (SCEPTRE), as a tool to quantify the fluorescence signal of immunolabeled histone marks or proteins structures at individual FISH-labeled genomic loci within individual cells (**fig. 1**).

**Figure 1.**
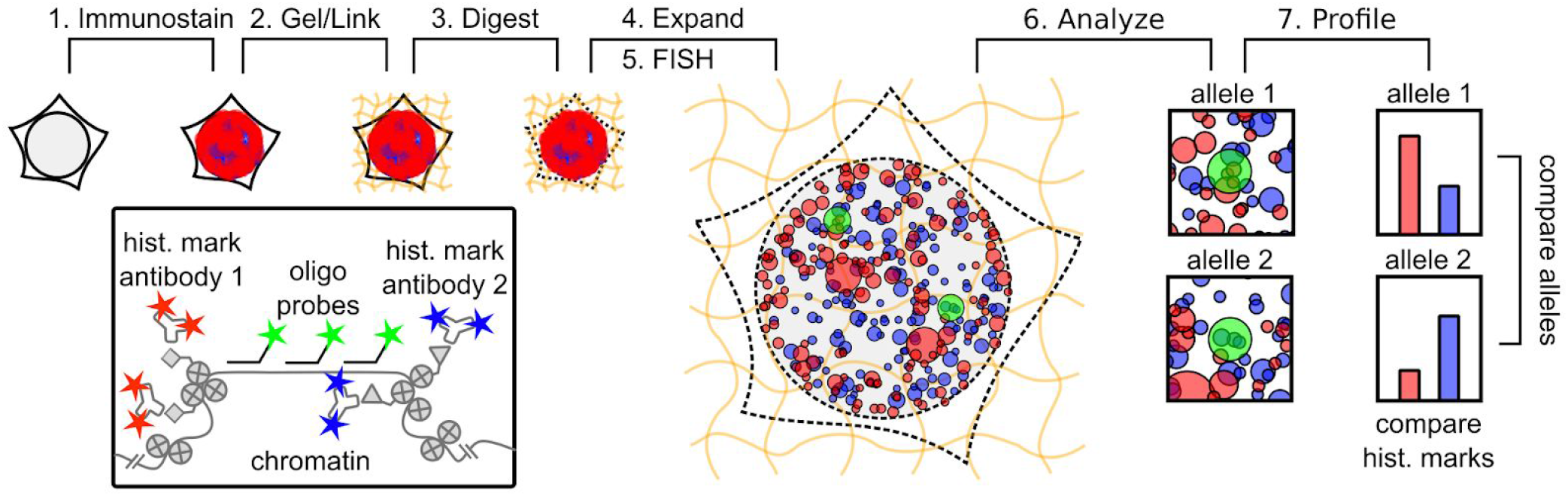
Workflow of SCEPTRE. **(1)** Histone marks or other protein structures are antibody-labeled in fixed cells. **(2)** The sample and antibodies are linked to a swellable hydrogel grown within the sample. **(3)** The sample is digested by proteinase K. **(4)** The hydrogel is expanded in water. **(5)** DNA loci, alleles from the same or different genes, are labeled by FISH. **(6)** The sample is imaged and relevant features are extracted for analysis. **(7)** An epigenetic profile is constructed for each cell, comparing histone mark levels between alleles or genes.

To test the ability of SCEPTRE to report on DNA-protein associations within the nucleus, we immunostained centromere-associated proteins while using DNA FISH to co-stain the repetitive alpha-satellite DNA of human centromeres (**fig. 2A**). ExM images revealed discrete regions corresponding to alpha-centromeres and centromere-associated proteins, as well as significant overlap between these two regions. To quantify this degree of overlap, we created an automated image analysis software routine in MATLAB to segment individual regions corresponding to centromeres and centromere-associated proteins, and then quantified their degree of co-localization (**sup. fig. 2**). From this analysis, we found that DNA-labeled regions had almost complete fractional overlap with centromere-associated proteins (0.97 ± 0.06). Furthermore, the distance between the nearest-neighbor centroids of the protein and DNA labeled regions was small relative to the average radius for either region (77 ± 85 nm versus 234 ± 68 nm (anti-cen.) and 224 ± 65 nm (α-cen.), respectively). We then quantified the fluorescence signal of labeled centromere-associated proteins at individual centromeric DNA clusters, along with randomly selected regions of comparable size to these FISH clusters (**fig. 2B**). While immunofluorescence and FISH-labeled regions maintained similar anti-centromere fluorescence signals, these regions showed much higher signals compared to randomly selected regions. Therefore, due to the high overlap between the FISH-labeled and immunolabeled regions, the high anti-centromere signal in the FISH-labeled regions, and the fact that the anti-centromere labeled structures did not shift dramatically between pre- and post-expansion (**sup. fig. 1**), we concluded that ExM can co-localize the signal of protein and DNA components of a genomic region within a nucleus.

**Figure 2.**
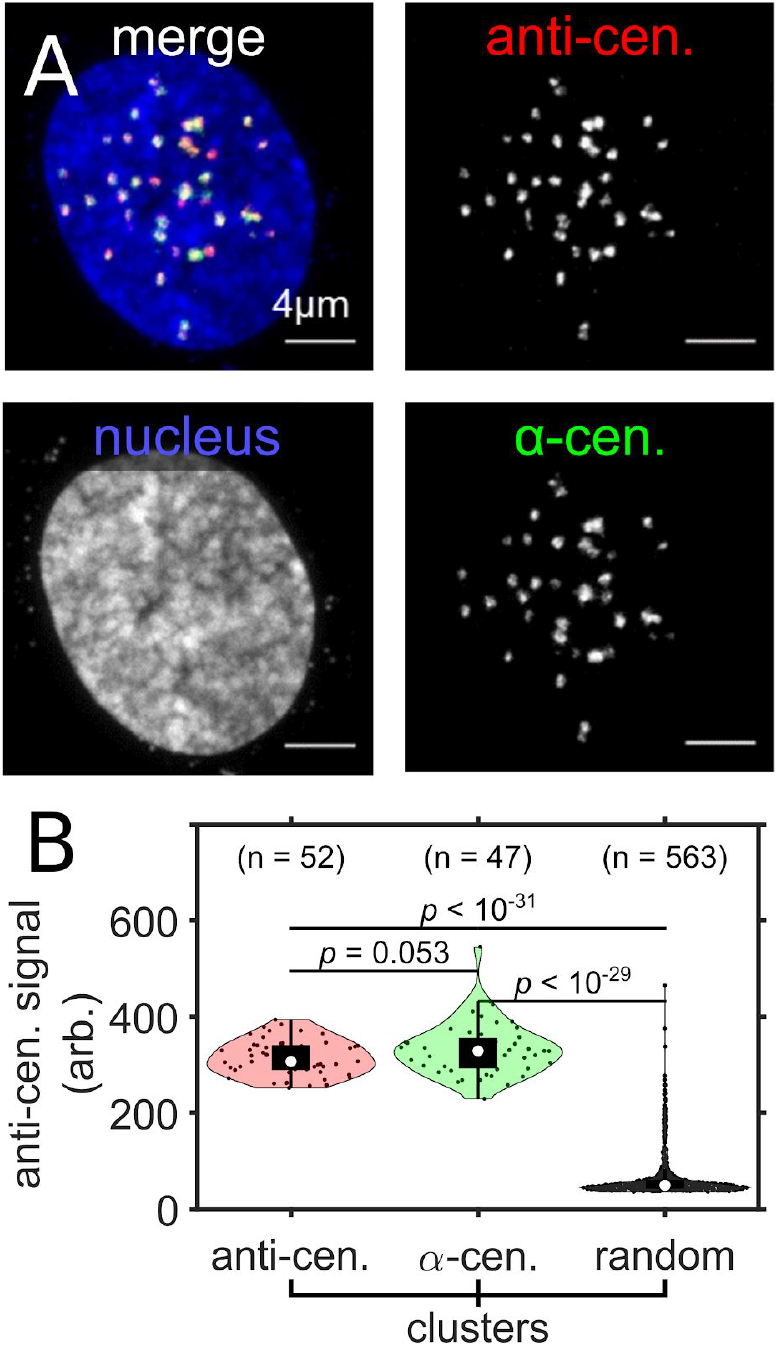
ExM reveals colocalization between centromere-associated proteins with repetitive centromeric DNA. **(A)** Maximum intensity projection image of an entire expanded RPE1 cell nucleus with immunolabeled centromere associated proteins (anti-cen., red), FISH labeled alpha-satellite DNA of centromeres (α-cen., green) and Hoechst-stained nucleus (blue). **(B)** the distribution of anti-centromere fluorescence signal (arb. = arbitrary units) in anti-centromere, α-centromere and in randomly selected region (random) clusters within the nucleus of the cell in **A.** Significance determined by a right-tailed Wilcoxon rank-sum test for anti-centromere against random, α-centromere against random, and α-centromere against anti-centromere cluster distributions. All scale bars are in pre-expansion units.

### SCEPTRE resolves multiple histone modifications at single gene loci in single cells

To determine whether SCEPTRE can distinguish between multiple histone marks at a single, non-repetitive genomic region, we concurrently visualized two histone marks, H3K4me3 and H3K27me3, at the house-keeping gene *GAPDH* (**fig. 3**). *GAPDH* encodes for glyceraldehyde-3-phosphate dehydrogenase, which is highly expressed in many cell types^42^ due to its essential role in metabolism;^43^ therefore, histone H3K4me3, commonly found at active gene promoters,^44^ is expected to be present at *GAPDH,* whereas histone H3K27me3, which is associated with repressed regions,^45^ is expected to be absent. Using SCEPTRE, we measured the fluorescence signals of immunolabeled H3K4me3 and H3K27me3 marks at the FISH-labeled *GAPDH* locus, along with the fraction of overlap between *GAPDH* and H3K4me3 or H3K27me3 clusters.

**Figure 3.**
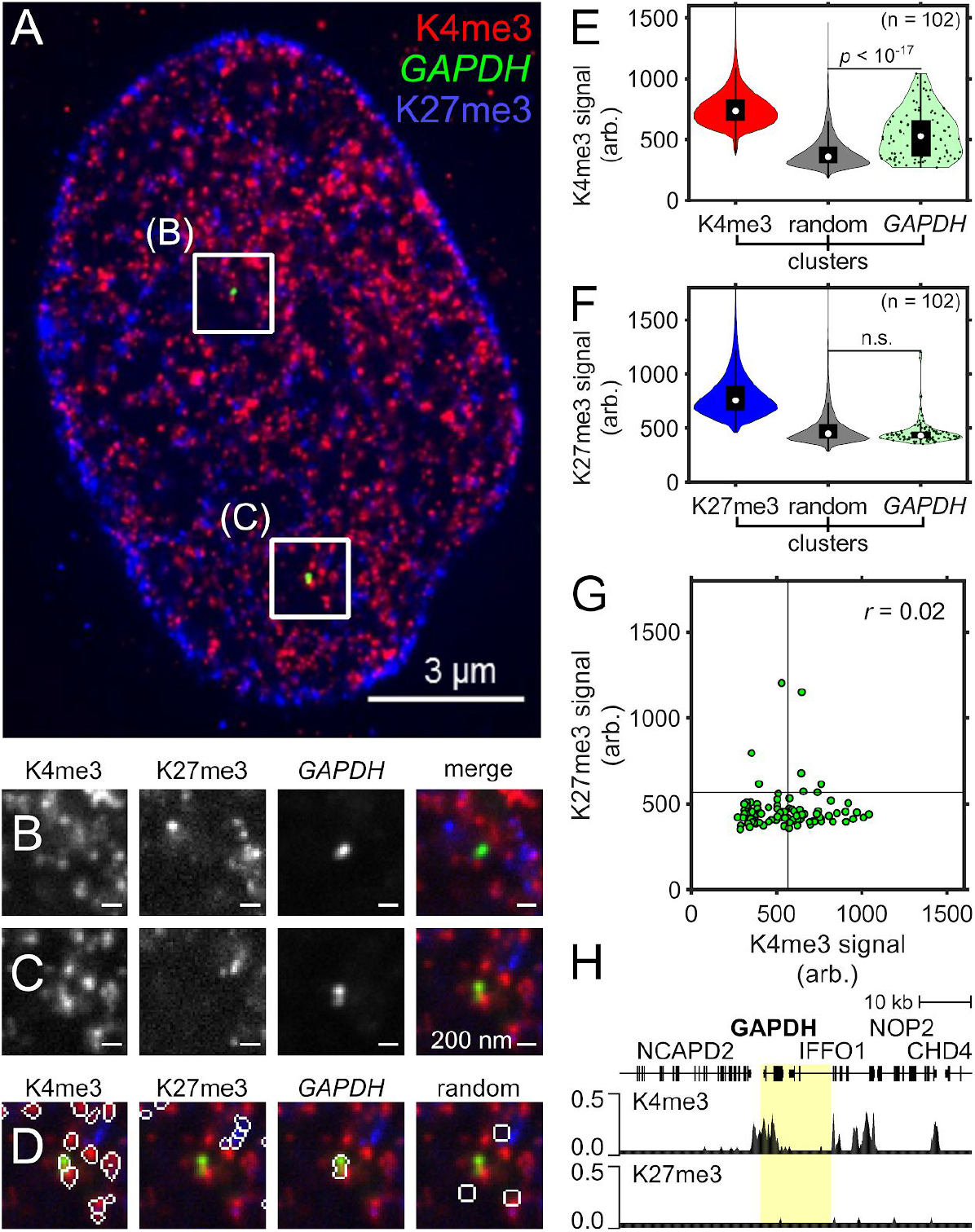
SCEPTRE distinguishes two histone marks at one genomic region. **(A)** An expanded RPE1 cell with immunolabeled H3K4me3 marks (K4me3, red) and H3K27me3 marks (K27me3, blue), and FISH-labeled *GAPDH* (green). **(B-C)** Zoomed in views of the approximate center plane of an image stack for each *GAPDH* allele in the cell seen in **A. (D)** Outline of the segmented regions for H3K4me3, *GAPDH,* H3K27me3 and randomly selected region clusters for the image plane seen in **C**. **(E)** Distribution of H3K4me3 fluorescence signal (arb. = arbitrary units) within H3K4me3, randomly selected regions (random) and *GAPDH* clusters. **(F)** Distribution of H3K27me3 fluorescence signal within H3K27me3, randomly selected regions and *GAPDH* clusters. **(G)** H3K27me3 and H3K4me3 fluorescence signals within *GAPDH* clusters (green). Black lines represent the threshold “on” level for each fluorescence signal. The correlation coefficient (r) between fluorescence signals within *GAPDH* is shown in the top-right corner of the plot. **(H)** CUT&RUN normalized counts for H3K4me3 (top) and H3K27me3 (bottom) marks in RPE1 cells for the FISH-targeted *GAPDH* region (highlighted). Cluster numbers for **E.** and **F.** are K4me3 = 343334, K27me3 = 478825, random = 8322, *GAPDH* = 102. Significance determined by a right-tailed Wilcoxon rank-sum test of histone mark fluorescence signals in *GAPDH* against random cluster distributions. All scale bars are in pre-expansion units.

From this analysis, we observed that *GAPDH* had much higher H3K4me3 fluorescence signal compared to H3K27me3 signal (**fig. 3E-F**). To our surprise, The H3K4me3 signal found at *GAPDH* varied greatly between loci, with some loci having high signals while others a more baseline level (**fig. 3G**). These results were the same when only one of both histone marks was labeled and imaged at *GAPDH* (**sup. fig. 3**), or when a different set of antibodies was used to label H3K4me3 and H3K27me3 in RPE1 cells (**sup. fig. 4**). Interestingly, histone mark signals were uncorrelated between *GAPDH* alleles in the same cell (**sup. fig. 5A-B**), suggesting histone mark levels at alleles from the same gene are independently regulated. When comparing these fluorescence signals to those obtained from randomly selected regions across the nucleus, *GAPDH* shared lower H3K27me3 signals and much higher H3K4me3 signals than those found at random regions. Similar to the fluorescence signal results, the mean fraction of overlap of *GAPDH* with H3K4me3 clusters was higher than with H3K27me3 clusters (0.21 ± 0.21 vs. 0.045 ± 0.11, respectively). To corroborate these results, we measured the density of H3K4me3 and H3K27me3 marks across the RPE1 genome for an ensemble of cells using CUT&RUN followed by sequencing.^27,28^ Analysis of the CUT&RUN sequencing results revealed that a substantial presence of H3K4me3 marks was found at the targeted *GAPDH* region and only background levels of H3K27me3 marks were found for this same region (**fig 3H**), with the closest repressed region observed ~500 kb away. These results demonstrate that SCEPTRE can distinguish between the abundance of two histone modifications at individual non-repetitive genomic regions within a nucleus.

H3K4me3 and H3K27me3 are generally thought to mark distinct chromatin states, though they have been reported to colocalize to form ‘bivalent domains’ on genes primed for transcription.^46,47^ We therefore investigated the relationship between H3K4me3 and H3K27me3 clusters across the nuclei of the RPE1 cells. As previously observed,^18^ H3K27me3 clusters preferentially inhabited the periphery of the nucleus, whereas H3K4me3 clusters were more evenly distributed (**fig. 3A**). There was a low fraction of spatial overlap between H3K4me3 clusters and H3K27me3 clusters (0.079 ± 0.14 H3K4me3 with H3K27me3, 0.12 ± 0.16 H3K27me3 with H3K4me3). The H3K4me3 fluorescence signal in H3K27me3 clusters, as well as the H3K27me3 signal in H3K4me3 clusters, was substantially low, albeit higher than the distribution of random regions (**sup. fig. 5C-D**). We therefore plotted the frequency of H3K4me3 and H3K27me3 signals within each of the other’s histone mark’s clusters, along with these signals found in randomly selected regions (**sup. fig. 6**). These plots show that H3K4me3 and H3K27me3 form largely non-overlapping clusters, though there exists a small fraction of clusters having high signal from both histone marks. These results suggest that H3K4me3 and H3K27me3 mostly form disjoint clusters, though a very small fraction may colocalize, consistent with the small percentage of nucleosomes that are found to harbor both marks at the same time.^48^

### SCEPTRE quantifies histone modification levels at multiple genomic loci in single cells

To test whether SCEPTRE can quantify histone mark signals at multiple genomic loci within the same cell, we designed a library of FISH probes to label three different genomic regions in RPE1 cells. The first region contains *MYL6,* a gene on Chr12 encoding myosin light chain-6 that is actively expressed in the eye;^49^ the second contains the *HOXC* gene cluster, which is normally active in progenitors but repressed upon differentiation;^45,50^ the third covers an internal region of long intergenic non-coding P53 induced transcript (*LINC-PINT*) variant, a non-coding transcript that is found on Chr7, which is broadly expressed across multiple tissues (**fig. 4**).^51^ As expected, bulk analysis of histone modifications at these loci using CUT&RUN revealed H3K4me3 peaks at *MYL6* and *LINC-PINT,* but not within the *HOXC* cluster. H3K27me3 marks, on the other hand, covered a large region encompassing the *HOXC* cluster, but were largely absent from *MYL6* and *LINC-PINT* (**fig. 4E**).

**Figure 4.**
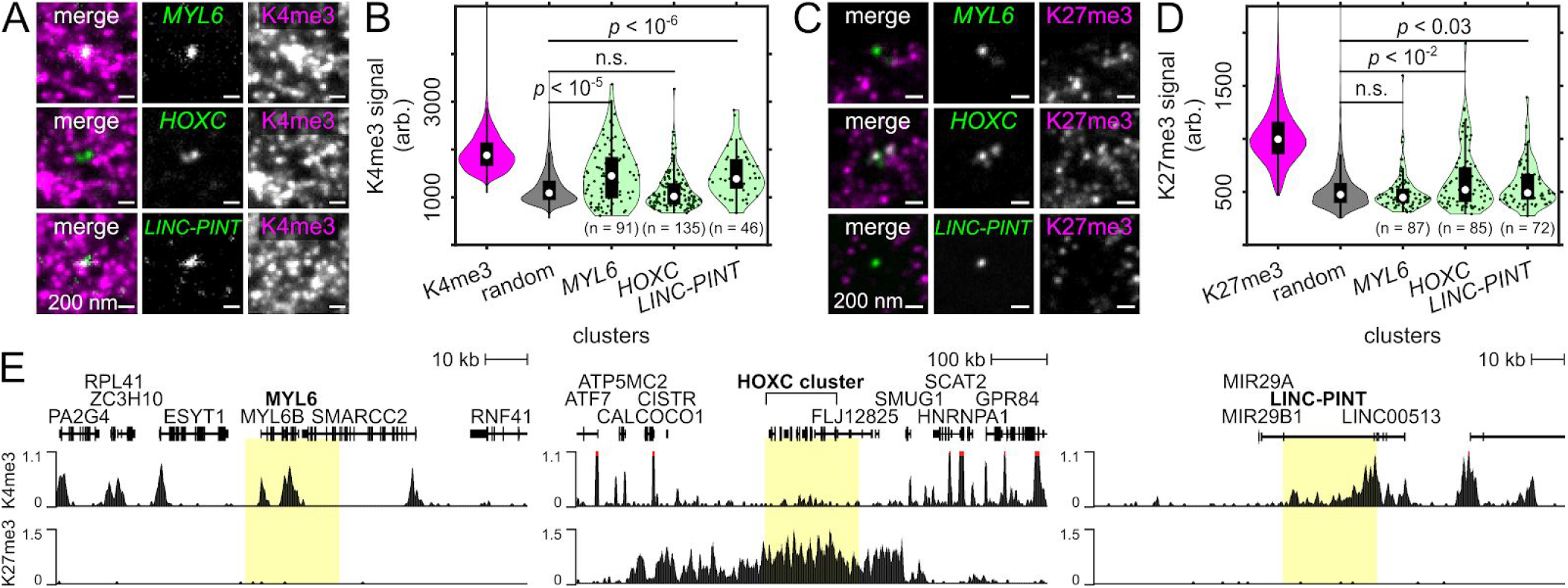
SCEPTRE quantifies one of two histone marks at three genomic regions. **(A)** Example image of the approximate center plane for each image stack of *MYL6, HOXC* or *LINC-PINT* FISH-labeled loci (green) from the same image stack of an expanded RPE1 cell immunolabeled for H3K4me3 marks (K4me3, magenta). **(B)** Distribution of H3K4me3 fluorescence signals (arb. = arbitrary units) within H3K4me3, randomly selected regions (random), *MYL6, HOXC* and *LINC-PINT* clusters (cluster numbers are K4me3 = 390331, random = 7421, *MYL6* = 91, *HOXC* = 135, *LINC-PINT* = 46). **(C)** Example image of the approximate center plane for each image stack of *MYL6*, *HOXC* or *LINC-PINT* FISH-labeled loci (green) from the same expanded RPE1 cell immunolabeled for H3K27me3 marks (K27me3, magenta). **(D)** Distribution of H3K27me3 fluorescence signals within H3K27me3, randomly selected regions, *MYL6, HOXC* and *LINC-PINT* clusters (cluster numbers are K27me3 = 196798, random = 6041, *MYL6* = 87, *HOXC* = 85, *LINC-PINT* = 72). **(E)** CUT&RUN normalized counts for H3K4me3 (top) or H3K27me3 (bottom) at the FISH-labeled *MYL6, HOXC* or *LINC-PINT* regions (highlighted). Significance determined by a right-tailed Wilcoxon rank-sum test of fluorescence signals in each FISH-labeled set against the random cluster distribution. All scale bars are in pre-expansion units.

In agreement with population-level results, H3K4me3 fluorescence signals measured using SCEPTRE were significantly higher at *MYL6* and *LINC-PINT* than at randomly chosen clusters (**fig. 4B,** *p*<10^-5^, *MYL6; p*<10^-6^, *LINC-PINT),* indicating enrichment of this histone modification at these two loci. Interestingly, both genes showed high H3K4me3 variability between individual loci, similar to what was observed at *GAPDH* loci. In contrast, H3K4me3 signals at the *HOXC* cluster were not significantly higher than those found at random regions. Consistently, H3K4me3 clusters showed greater overlap with the *MYL6* and *LINC-PINT* loci than at the *HOXC* cluster, where it appeared to be visibly excluded (0.23 ± 0.26, *MYL6* and 0.20 ± 0.24, *LINC-PINT* vs 0.068 ± 0.16, *HOXC*). Therefore, SCEPTRE detects differences in H3K4me3 levels between multiple genomic regions seen with population-level analysis, agreeing with the results obtained by CUT&RUN. We note that H3K4me3 mark signals were largely uncorrelated between two alleles of each gene in each cell (**sup. fig. 7A**), as well as between the loci of different genes in the same cell **(sup. fig. 8A)**, indicating that the levels of this histone mark are largely independent across loci within individual cells.

Also in agreement with bulk analysis, The *HOXC* cluster showed higher H3K27me3 fluorescence signal compared to random clusters, and higher H3K27me3 signal compared to *MYL6* or *LINC-PINT* (**fig. 4D**). Strikingly, H3K27me3 levels varied substantially between different HOXC clusters, with a substantial fraction of *HOXC* clusters with either low or even baseline levels of H3K27me3. Although *MYL6* did not have significantly higher H3K27me3 fluorescence levels compared to random regions, *LINC-PINT* did, despite an absence of H3K27me3 marking seen in CUT&RUN data. The presence of H3K27me3 at some *LINC-PINT* loci may reflect the looping of this locus to a different genomic region where H3K27me3 is present; to investigate this possibility, we consulted a previously published HiC data set for the RPE1 cell line;^52^ indeed *LINC-PINT* maintained high frequency contacts with an adjacent H3K27me3 domain (**sup. fig. 9**). These results suggest that, at the current spatial resolution, adjacent genomic regions can influence each other’s histone mark levels detected by SCEPTRE. That being said, the SCEPTRE results broadly agree with the results obtained by CUT&RUN and can distinguish between the chromatin modification states of multiple genes in the same cell (e.g., *MYL6* and *HOXC*). As seen with the H3K4me3 marks at these genomic regions, we observed no relationship between the H3K27me3 levels for two alleles of the same gene (**sup. fig. 7B**) or for alleles from different genes (**sup. fig. 8B**) in the same cell.

### H3K4me3 modifications coincide with paused RNA polymerase II at a transcriptionally active locus

H3K4me3 levels have been reported to correlate with active transcription^53^ and a model has been proposed where H3K4me3 facilitates the loading of RNA polymerase II, which remains paused proximally to the gene’s promoter until a subsequent release step.^54^ However, this model was based on separate population-level measurements of H3K4me3 and RNA polymerase II, and did not distinguish whether both components coincide directly at the same time at single loci in cells. To test whether both H3K4me3 and paused RNA polymerase II were present simultaneously at *GAPDH,* we performed SCEPTRE with H3K4me3 and the post-translationally modified form of paused RNA polymerase II during transcription initiation, where the Serine 5 of the repeat C-terminal domain of RNA polymerase II is phosphorylated (**fig. 5**).^55–57^

**Figure 5.**
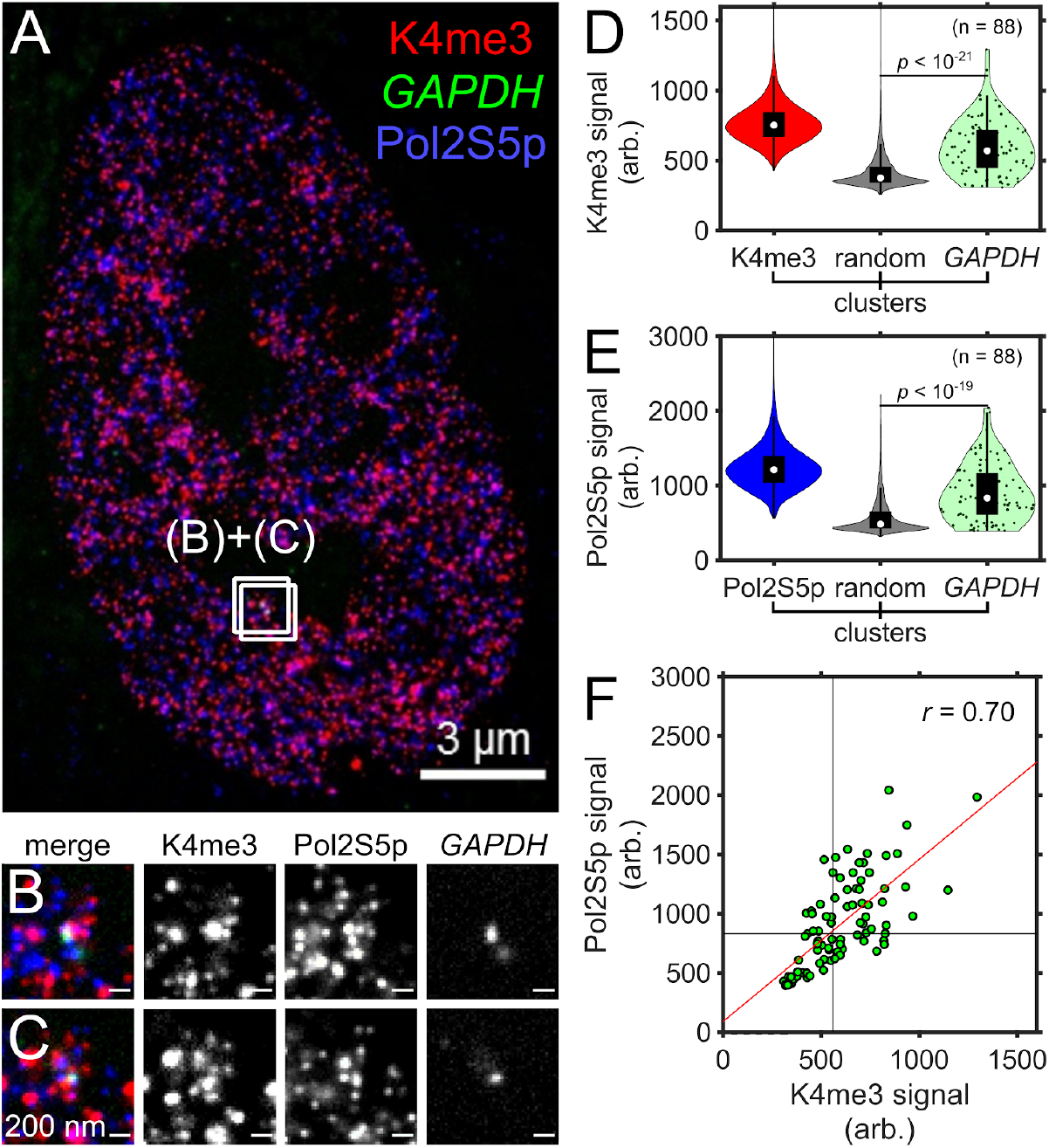
SCEPTRE compares H3K4me3 and paused RNA polymerase II signals at a single genomic region. **(A)** An expanded RPE1 cell with immunolabeled H3K4me3 marks (K4me3, red) and paused RNA polymerase II (Pol2S5p, blue), and FISH-labeled *GAPDH* (green). **(B-C)** Zoomed in views of the approximate center plane of an image stack for each *GAPDH* allele in the cell seen in **A. (D)** Distributions of H3K4me3 fluorescence signal (arb. = arbitrary units) within H3K4me3, randomly selected regions (random) and *GAPDH* clusters. **(E)** Distribution of paused RNA polymerase II fluorescence signal within paused RNA polymerase II, randomly selected regions and *GAPDH* clusters. **(F)** H3K4me3 and paused RNA polymerase II fluorescence signals within *GAPDH* clusters (green). Black lines represent the threshold “on” level for each fluorescence signal, while the red line represents the linear regression. The correlation coefficient (r) between fluorescence signals within *GAPDH* is shown in the top-right corner of the plot. Cluster numbers in **D.** and **E.** are K4me3 = 440298, Pol2S5p = 542245, random = 8240, *GAPDH* = 88. Significance determined by a right-tailed Wilcoxon rank-sum test of histone mark fluorescence signals in *GAPDH* against random cluster distributions. All scale bars are in pre-expansion units.

We detected a large coincidence between H3K4me3 and paused RNA polymerase II, both at the *GAPDH* locus and also more broadly in the nucleus. At individual *GAPDH* loci, there were high signals from both H3K4me3 and paused RNA polymerase II (**fig. 5B-E**), such that there was also a strong correlation between these signals (**fig. 5F**, *r* = 0.70). Consistently, *GAPDH* overlapped with both H3K4me3 and paused RNA polymerase II clusters (0.23 ± 0.21 and 0.21 ± 0.19, respectively). Similarly to H3K4me3, paused RNA polymerase II signals were uncorrelated between *GAPDH* loci in the same cell (**sup. fig. 10**). In the nucleus more broadly, there was also substantial colocalization between H3K4me3 clusters and paused RNA polymerase II clusters (**fig. 5B,** fraction of overlap 0.19 ± 0.21), as well as a strong correlation between these two signals in randomly selected region clusters (**sup. fig. 11C**, *r*= 0.68).

In contrast, no correlation was seen at *GAPDH* between H3K4me3 and H3K27me3 signals (**fig. 3G**, *r* = 0.02), or between H3K27me3 and paused RNA polymerase II (**sup. fig. 12F**, *r* = 0.04). On a broader level, there was also little to no correlation in random regions between H3K4me3 and H3K27me3 (**sup. fig. 5C,** *r* = 0.18), or between H3K27me3 and paused RNA polymerase II (**sup. fig. 13C**, *r* = 0.17). Interestingly, when H3K27ac, another active histone mark, was concurrently visualized with paused RNA polymerase II, some correlation was seen between these two signals, with *r*= 0.43 at *GAPDH* (**sup. fig. 14F**), and *r* = 0.59 at random regions (**sup. fig 15C**). However, the fraction of *GAPDH* loci with high H3K27ac signals were smaller compared to that with high paused RNA polymerase II signals, suggesting that H3K27ac and the phosphorylation indicative of paused RNA polymerase II play distinct roles in the transcriptional cycle. Together, these results are consistent with a close regulatory relationship between H3K4me3 modifications and the loading of paused RNA polymerase II, both at *GAPDH* and more broadly across the nucleus.

## Discussion

SCEPTRE is a new method capable of profiling chromatin states at multiple genomic loci within the 3D nuclear context of a cell by combining immunofluorescence with DNA *in situ* labeling by means of ExM. This combination provides rapid acquisition of histone mark fluorescence signals at a resolution of ~75 nm, sufficient to quantify histone mark abundance at individual genomic loci. In contrast to sequencing-based methods, SCEPTRE provides quantitative measurements of physical properties, such as overlap, density, and position within the nucleus for more than one histone mark at multiple genomic regions. Such measurements reveal a heterogeneity in chromatin states that has been previously masked in ensemble sequencing-based methods.

There are limitations to SCEPTRE compared to other histone mark profiling methods. Sequencing based methods can achieve nucleosome level resolution for histone mark mapping across an entire genome, such as in the case of CUT&RUN.^27^ Since SCEPTRE relies on DNA FISH, detection of genes by *in situ* labeling often is limited to a minimum labeling size of over 10 kb, since smaller regions are detected with lower efficiency. However, genome organization is thought to occur at a larger scale than that of the single nucleosome. Nucleosomes are known to organize as clusters throughout the nucleus, with spatial sizes ranging around 50-100 nm,^17^ a size that corresponds to roughly 10 kb of genomic DNA, depending on the region’s activity state.^19^ The scale increases further when observing Topologically Associating Domains (TADs), which are genomic regions of around 200 kb to 1 Mb in size that maintain similar epigenetic and regulatory landscapes,^58,59^ or the smaller sub-TADs that are ~185 kb,^60^ with spatial sizes of ~160 nm.^61^ Even larger than 1 Mb are chromatin A and B compartments who are associated with broader open (active) and closed (repressed) states,^62^ with spatial sizes on the μm scale.^63,64^ Since SCEPTRE has allowed us to profile multiple genes at the lower scale of this genomic organization, there is potential to build upon this technique in order to target larger genomic regions by using multiplex FISH methods, such as MERFISH,^65^ seqFISH,^66^ ORCA^67^ or OligoFISSEQ.^68^ These methods would allow SCEPTRE to interrogate the relationships between histone modifications and gene activity at a variety of developmentally-regulated genes, and at increasingly larger scales of genome organization.

SCEPTRE revealed heterogeneity in the levels of H3K4me3 at active genomic regions such as *GAPDH, MYL6* and *LINC-PINT.* This variability suggests active genes do not maintain steady histone mark levels, but can instead switch between different modification states in a dynamic manner. This heterogeneity may account for the low frequency of histone mark detection at individual sites in sequencing-based single cell methods, such as scChIP-seq and scCUT&TAG, where only a fraction of reads fall within known domains for a given histone mark, and only a fraction of known domains have reads within them.^9,69^ However, because SCEPTRE revealed a close relationship between H3K4me3 and paused RNA polymerase II at individual loci, this heterogeneity may not simply reflect detection limitations, but may be closely related to the transcriptional state of each gene. Given that genes are transcribed in bursts,^70^ where polymerase recruitment happens intermittently,^71^ it is plausible that H3K4me3 marks and the phosphorylation indicative of paused RNA polymerase II are dynamically added during a transcriptional burst, but later removed at a later stage in the transcriptional cycle. Moving forward, it would be useful to utilize SCEPTRE to further visualize H3K4me3 and other histone marks alongside different stages of transcription, to elucidate how histone marks participate in the regulation of gene transcription.

Although H3K27me3 marks were enriched in the repressed *HOXC* cluster compared to active genes, consistent with population-level measurements, H3K27me3 levels at this cluster also showed striking heterogeneity between individual loci, similar to that observed for H3K4me3. The apparent absence of H3K27me3 marks on some *HOXC* loci may reflect technical limitations, such as inefficient or non-specific antibody labeling, or other unrecognized technical variability. Alternatively, these results may reflect true, and therefore under-appreciated heterogeneity in H3K27me3 levels at repressed gene loci. Gene repression at the Hox gene cluster requires PRC1, a protein complex that mediates chromatin compaction and gene silencing.^72^ PRC1 binds to H3K27me3, an interaction that explains the co-localization of these two factors in the genome; however, PRC1 can also bind genomic loci independently of H3K27me3, and thus stably maintain a repressed state at the *HOXC* locus, even when H3K27me3 is absent.^73^ In such a picture, H3K27me3 levels may fluctuate at individual loci, and reach baseline levels without loss of PRC1 binding and stable gene repression. To investigate this further, it will be helpful to use SCEPTRE to interrogate the composition of polycomb domains throughout the genome, as well as with other methods that can visualize chromatin state dynamics in living cells.

Lastly, there are certain factors that influence the way SCEPTRE profiles the epigenetic state of genes. As demonstrated in the example of the *LINC-PINT* region (**fig. 4** and **sup. fig 9**), the 3D context of a region can influence its epigenetic profile. This “crosstalk” from neighboring regions is most-likely due to the fact that genes with different epigenetic states may be found closer than the resolution of ExM provides at a 4× expansion factor (~75 nm). If so, methods that achieve better resolution, such as iterative ExM^74^ or by combining ExM with SIM,^75^ would provide a greater distinction between epigenetic states of neighboring genes while using SCEPTRE. Another factor that may play a role in profiling are the genetic elements within a targeted region. The labeled *LINC-PINT* region was an internal sequence of a gene, which showed a different distribution of H3K4me3 signals compared to the *MYL6* region, whose promoter was found at the center of the labeled region. Therefore, considering the 3D-context of chromatin within cells (seen by HiC from a previous study)^52^ and the epigenetic landscape of a genomic sequence (seen by CUT&RUN in this study) can help with either selecting each targeted region for SCEPTRE, or in determining the epigenetic state for each region within a cell. Further improvements in multiplexing and resolution would allow SCEPTRE to systematically profile chromatin states in the genome, providing new insights into our understanding of genome structure and function.

## Acknowledgements

This work is supported by the University of Washington (H.Y.K. and J.C.V.), the Chan Zuckerberg Initiative (H.Y.K and J.C.V.), a fellowship by the NHLBI F31HL142132 (M.A.W.) and a training grant by the NIGMS T32GM008268 (M.A.W). The authors would like to thank L. Worderman (University of Washington) for providing the hTERT RPE1 cell line; S. Henikoff (Fred Hutchinson Cancer Research Center) for the generous gifts of pA-MNase and spike-in yeast DNA; B. Beliveau (University of Washington) for assistance with probe design; and the Biology Imaging Facility at the University of Washington for providing imaging assistance.

## Contributions

M.A.W., K.K.H.N., H.Y.K. and J.C.V. designed the experiments. M.A.W., K.K.H.N., A.R.H., N.A.P. and P.H.B.N. contributed to the experimental setup. M.A.W. and K.K.H.N. obtained the results. M.A.W., K.K.H.N., A.R.H., N.A.P. and P.H.B.N. contributed to the results analysis. M.A.W., H.Y.K. and J.C.V wrote the paper and all authors commented on the manuscript. H.Y.K. and J.C.V. supervised the project.

---

**Supplementary figure 1.**
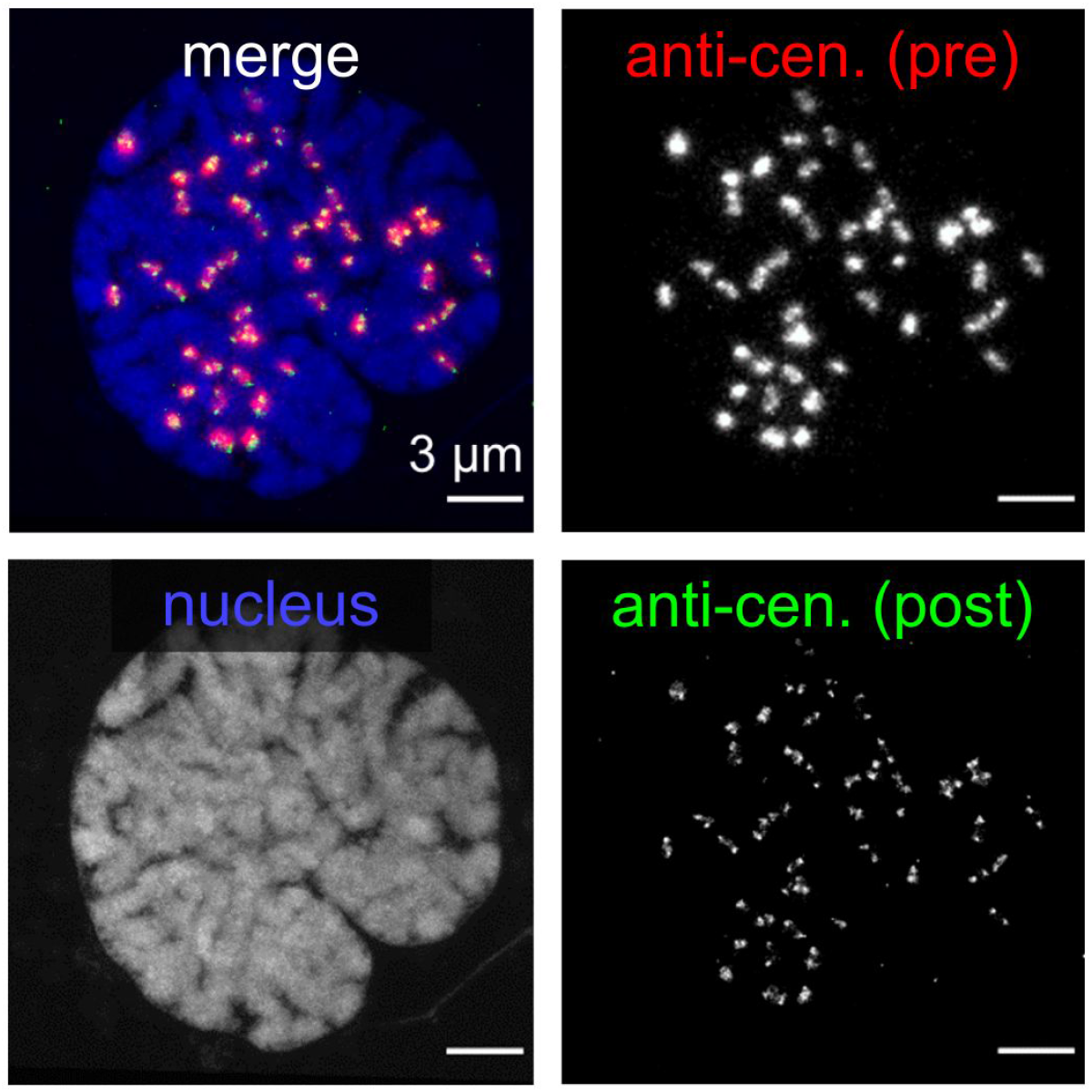
Correlative imaging of anti-centromere stain before and after expansion. Anti-centromere imaged post-expansion (post, green), is aligned by similarity transform to the same stain imaged pre-expansion (pre, red) and visualized in the context of the post-expansion nucleus labeled by Hoechst (blue). All scale bars are in pre-expansion units.

**Supplementary figure 2.**
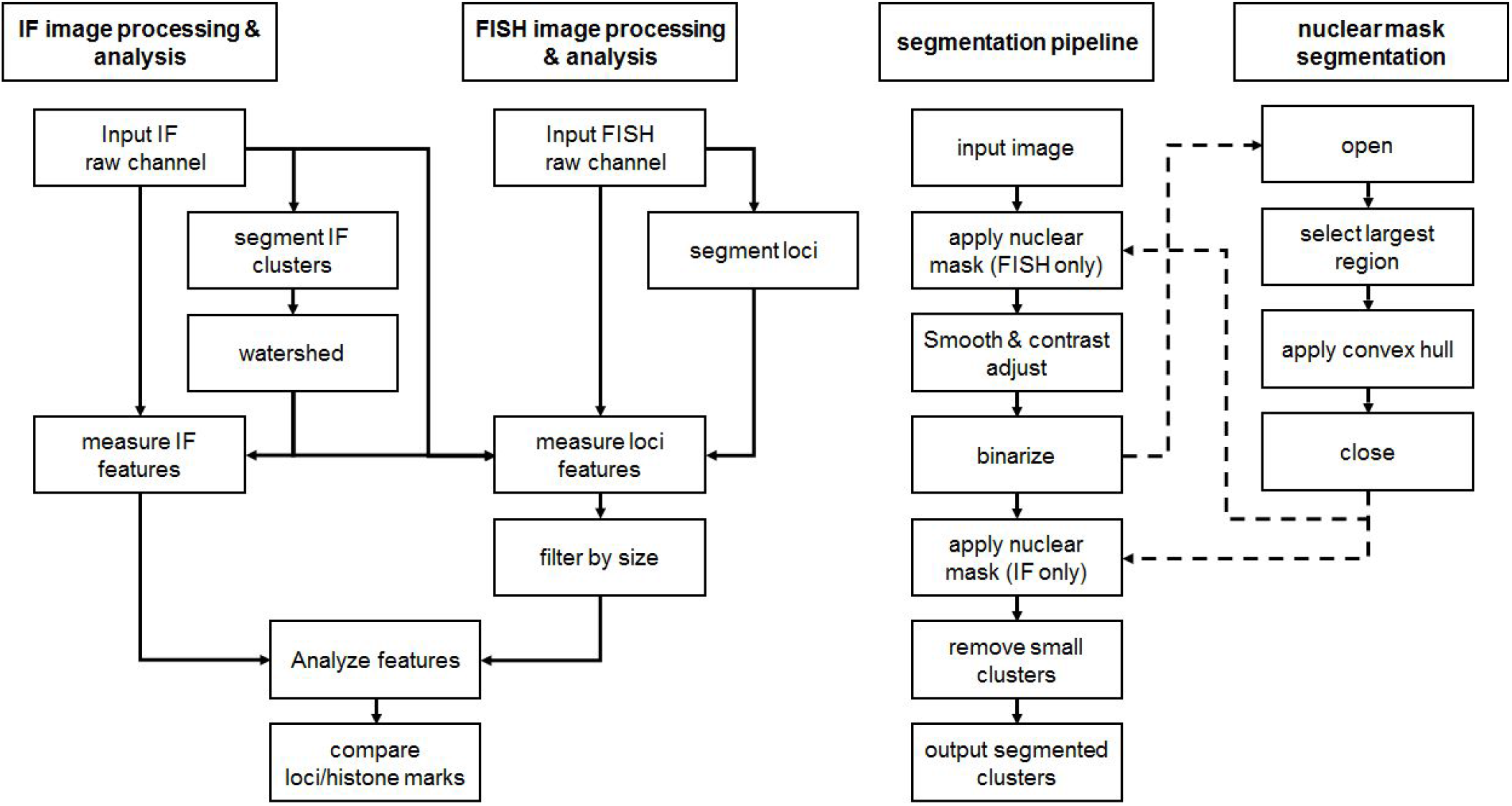
Image processing schematic for SCEPTRE. Raw images obtained from the immunofluorescence (IF) of protein structures are segmented with the following steps: smooth with a gaussian filter, then contrast adjust with an adaptively determined threshold per cell; binarize either by an Otsu method, or by a Laplace filter followed by selection of all negative values; apply a nuclear mask; watershed. After the segmentation of the nuclear channel and the immunofluorescence channels, the FISH raw channel is then segmented in the same manner with the following exceptions: a nuclear mask is applied before smoothing and contrast adjustment, and no watershed is applied. Features, including mean fluorescence intensity and fraction of overlap with segmented clusters from each immunofluorescence channel are identified for all segmented clusters within a channel. FISH clusters are further filtered by size. The nuclear mask is generated with the following additional segmentation steps: smooth and contrast adjust either a Hoechst stain channel or one of the present immunofluorescence channels; open image to fuse clusters within the nucleus; select largest region encompassing the nucleus; apply a convex hull; close segmented nucleus; apply to immunofluorescence and FISH channels (for more details, see Materials and Methods and **sup. table 2**).

**Supplementary figure 3.**
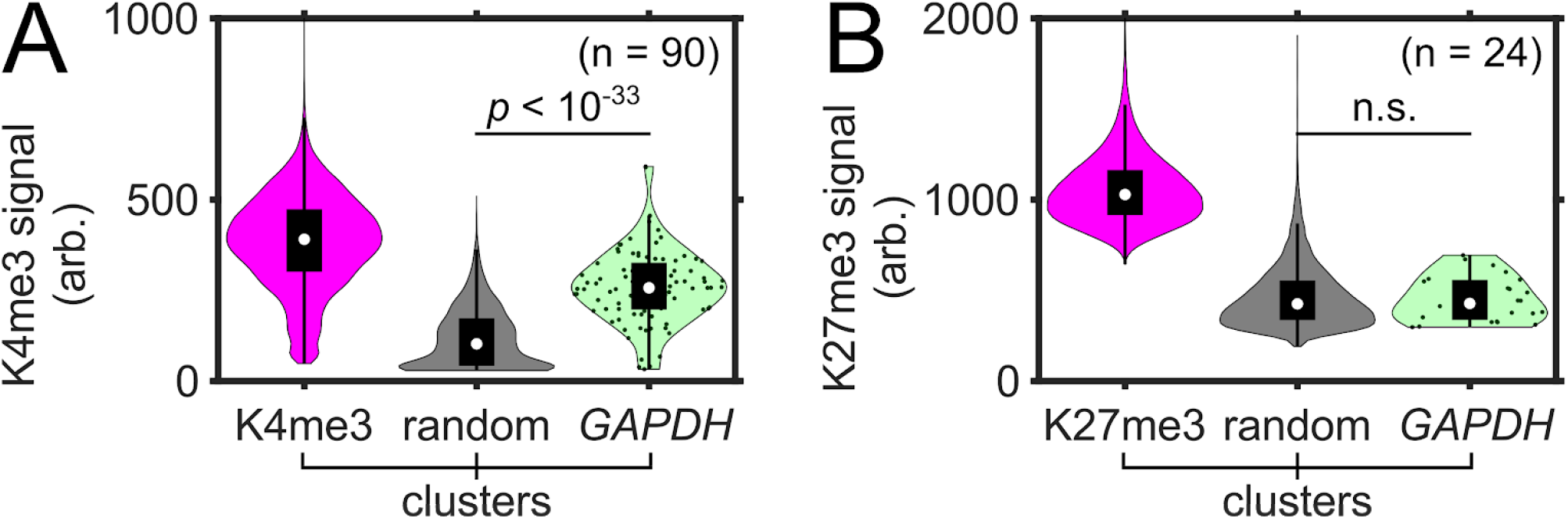
SCEPTRE measures signal of single-immunolabeled histone marks at *GAPDH* in RPE1 cells. **(A)** Distributions of H3K4me3 (K4me3) fluorescence signal (arb. = arbitrary units) within H3K4me3, randomly selected regions (random) and *GAPDH* clusters from single-immunolabeled expanded RPE1 cells. Cluster numbers are K4me3 = 196194, random = 5744, *GAPDH* = 90. **(B)** Distribution of H3K27me3 (K27me3) fluorescence signal within H3K27me3, randomly selected regions and *GAPDH* clusters from single-immunolabeled expanded RPE1 cells. Cluster numbers are K27me3 = 60235, random = 6504, *GAPDH* = 24. Significance determined by a right-tailed Wilcoxon rank-sum test of histone mark fluorescence signal in *GAPDH* against random cluster distributions.

**Supplementary Figure 4.**
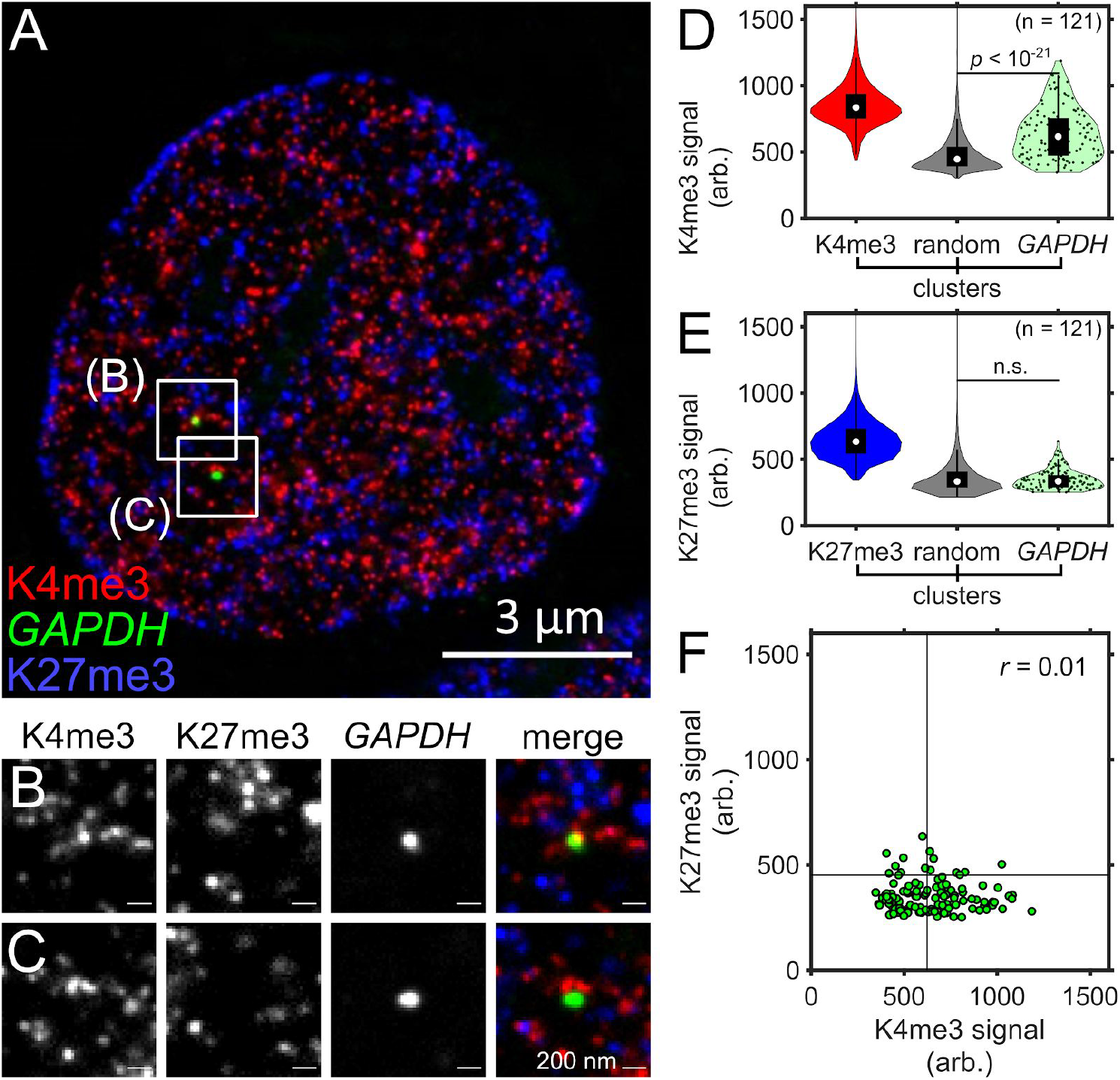
SCEPTRE shows reproducible results with a different set of antibodies. **(A)** An expanded RPE1 cell with immunolabeled H3K4me3 marks (K4me3, red) and H3K27me3 marks (K27me3, blue), and FISH-labeled *GAPDH* (green), using an alternative set of antibodies to **figure 3**. **(B-C)** Zoomed in views of the approximate center plane of an image stack for each *GAPDH* allele in the cell seen in **A. (D)** Distributions of H3K4me3 fluorescence signal (arb. = arbitrary units) within H3K4me3, randomly selected regions (random) and *GAPDH* clusters. **(E)** Distribution of H3K27me3 fluorescence signal within H3K27me3, randomly selected regions and *GAPDH* clusters. **(F)** H3K27me3 and H3K4me3 fluorescence signals within *GAPDH* clusters (green). Black lines represent the threshold “on” level for each fluorescence signal. Cluster numbers are K4me3 = 250644, K27me3 = 262307, random = 7406, *GAPDH* =121. The correlation coefficient (r) between fluorescence signals within *GAPDH* is shown in the top-right corner of the plot. Significance determined by a right-tailed Wilcoxon rank-sum test of histone mark fluorescence signals in *GAPDH* against random cluster distributions. All scale bars are in pre-expansion units.

**Supplementary figure 5.**
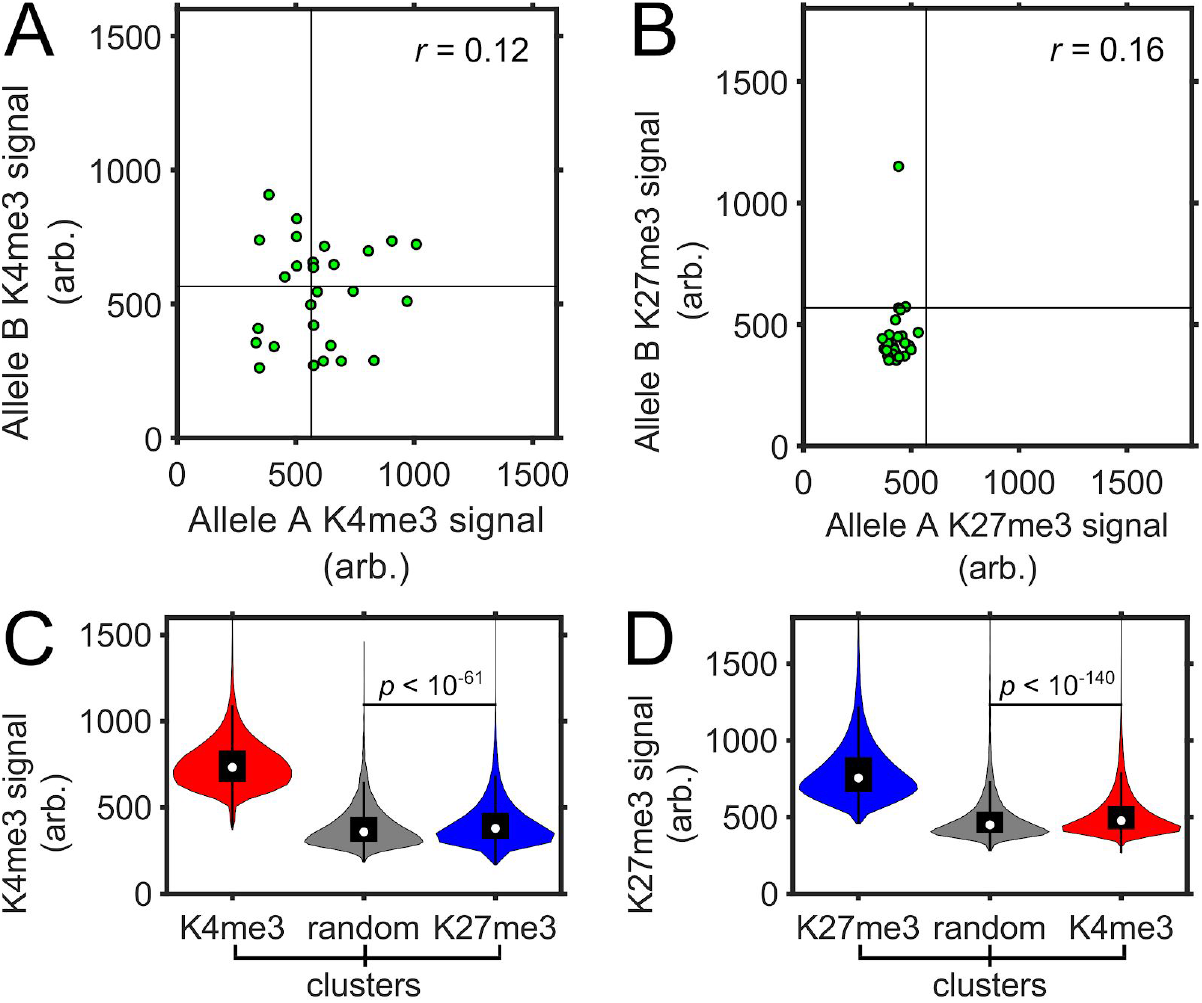
SCEPTRE compares H3K4me3 and H3K27me3 signals between different *GAPDH* alleles in the same cell, or between histone mark cluster distributions. **(A-B)** Fluorescence signal (arb. = arbitrary units) of either H3K4me3 (K4me3) in **A.**, or H3K27me3 (K27me3) in **B.**, in *GAPDH* alleles within the same cell from the data set in **figure 3** (one locus from each cell containing 2-4 loci is randomly assigned as allele A, and a second locus as allele B). Black lines represent the threshold “on” level for each histone mark fluorescence signal. The correlation coefficient (r) is shown on the top-right corner of each plot. **(C-D)** Fluorescence signal for either H3K4me3 in **C.**, or H3K27me3 in **D**., for each distribution of H3K4me3 (red), H3K27me3 (blue) and randomly selected region (random, gray) clusters within the cells in **figure 3**. Significance determined by a right-tailed Wilcoxon rank-sum test of fluorescence signals in each histone mark cluster set against random cluster distributions.

**Supplementary figure 6.**
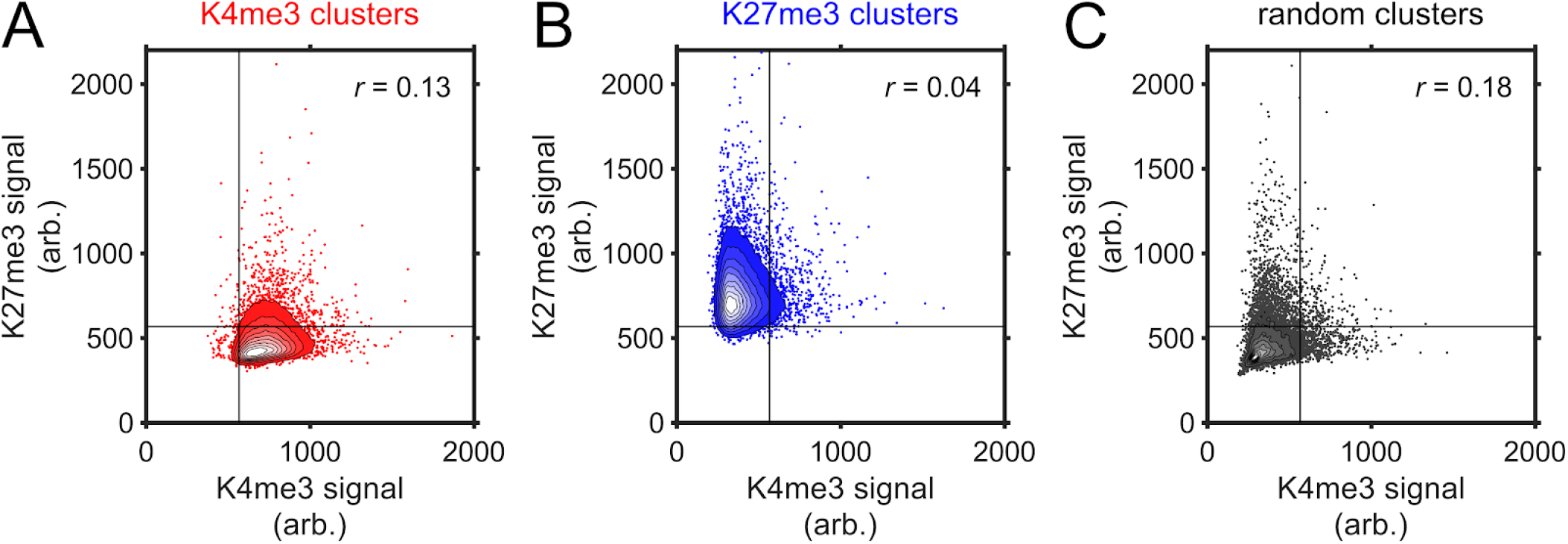
SCEPTRE compares H3K4me3 and H3K27me3 signals within segmented immunostained and random clusters. Contours for the fluorescence signal (arb.=arbitrary units) frequency of H3K4me3 (K4me3) and H3K27me3 (K27me3) in the cluster sets of H3K4me3 (red) in **A.**, H3K27me3 (blue) in **B.**, and randomly selected regions (random, gray) in **C**. Straight black lines represent the threshold “on” level for each fluorescence signal. Contours have uniformly spaced steps ranging from 0.1 to 0.9 frequency and represent all clusters obtained for cells in **figure 3**. The remaining scatter in **A.** and **B.** is a 100-fold downsample of the original data by random selection for plot representation purposes. Correlation coefficients (r) for each data set, which are calculated before downsampling, are shown in the top-right corner of each plot.

**Supplementary figure 7.**
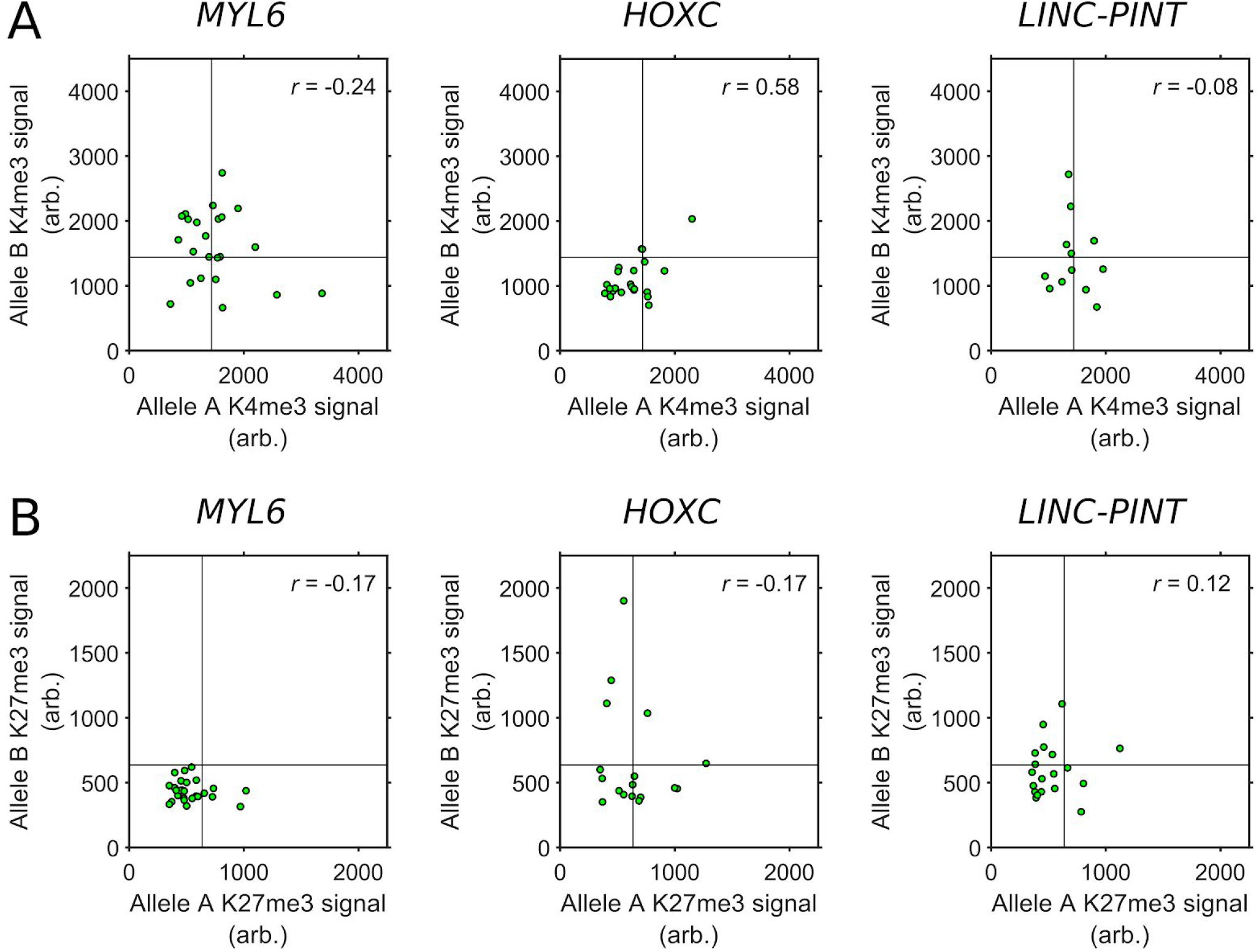
SCEPTRE compares H3K4me3 or H3K27me3 signals between alleles of one of multiple genes in the same cell. Fluorescence signal of either H3K4me3 (K4me3) in **A.**, or H3K27me3 (K27me3) in **B.**, in *MYL6, HOXC* or *LINC-PINT* alleles from the same cell (one locus from each cell containing 2-4 loci is randomly assigned as allele A, and another one as allele B). Black lines represent the threshold “on” level for each histone mark fluorescence signal. The correlation coefficient (r) for each set is shown in the top-right corner of each plot.

**Supplementary figure 8.**
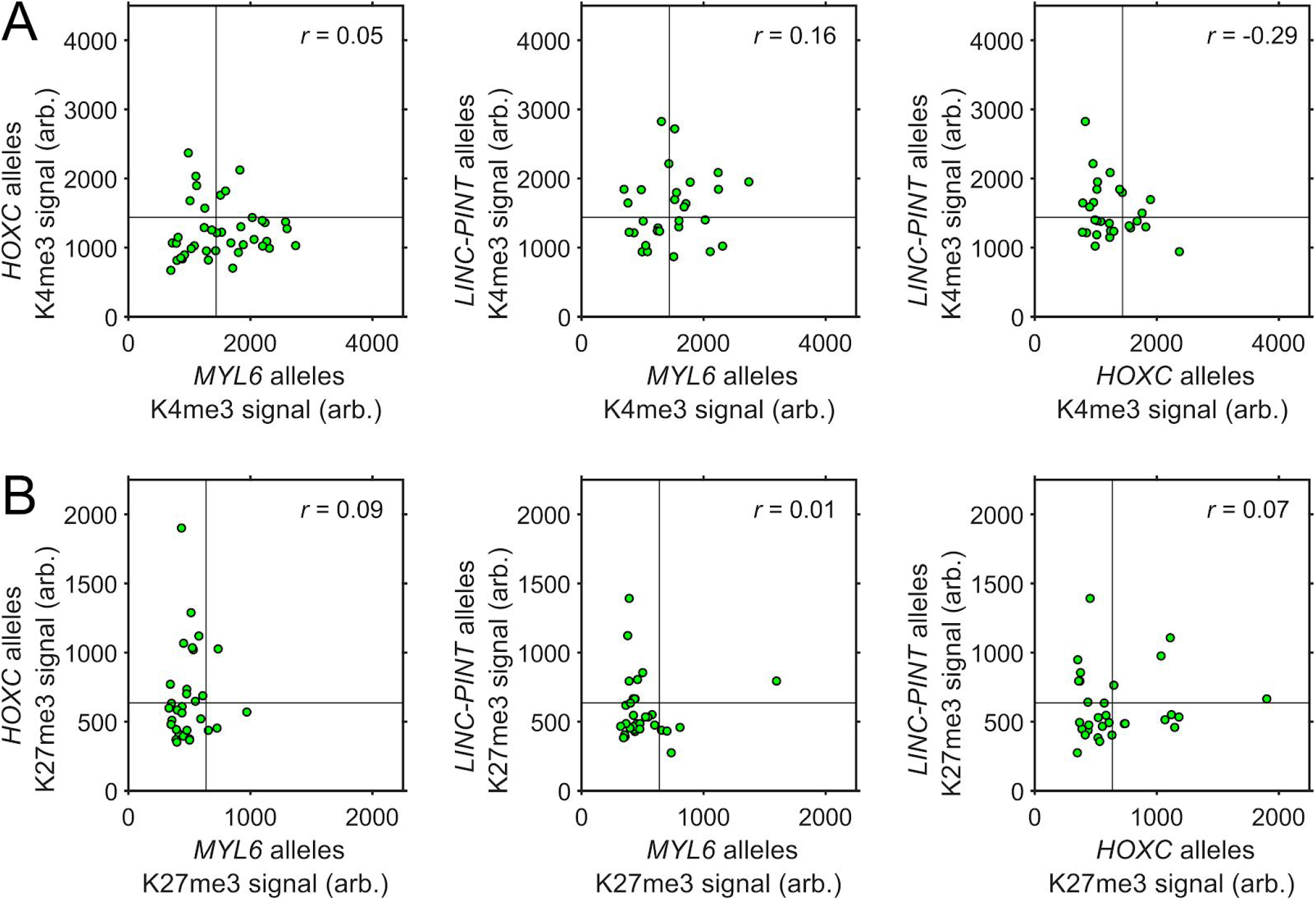
SCEPTRE compares H3K4me3 or H3K27me3 signals between alleles from different genes in the same cell. Comparison of the fluorescence signal (arb.= arbitrary units) of either H3K4me3 (K4me3) in **A.**, or H3K27me3 (K27me3) in **B.**, between randomly selected alleles of *MYL6, HOXC* and/or *LINC-PINT* within the same cell. Black lines represent the threshold “on” level for each histone mark fluorescence signal. The correlation coefficient (r) for each set is shown in the top-right corner of each plot.

**Supplementary Figure 9.**
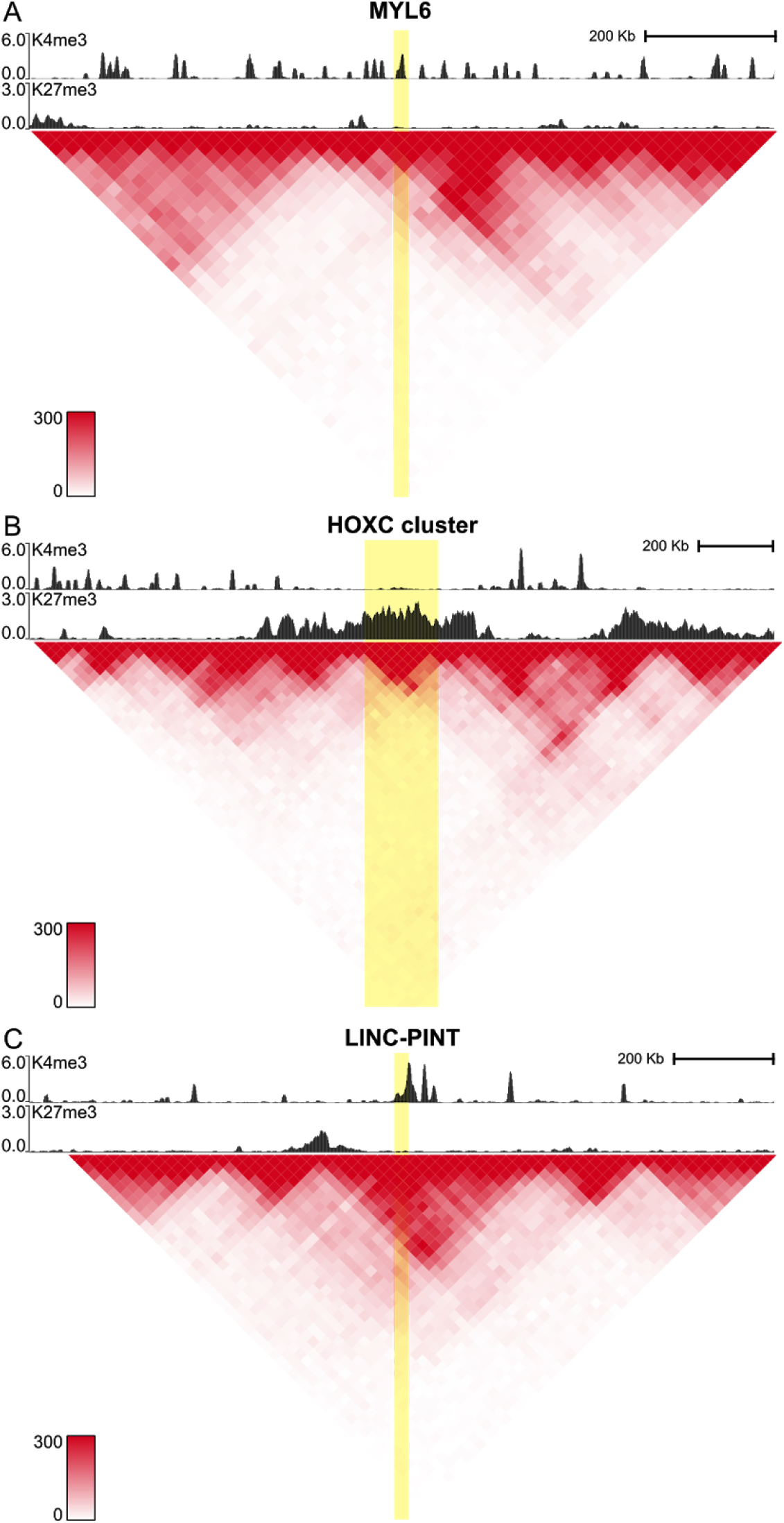
Analysis of Hi-C for targeted genomic regions in RPE 1 cells. Hi-C data, previously published,^52^ along with H3K4me3 (K4me3) and H3K27me3 (K27me3) CUT&RUN normalized counts for *MYL6* (**A**), *HOXC* (**B**) and *LINC-PINT* (**C**) targeted regions (highlighted). Heat map score between 0 - 300 reads in 25 kb bins.

**Supplementary figure 10.**
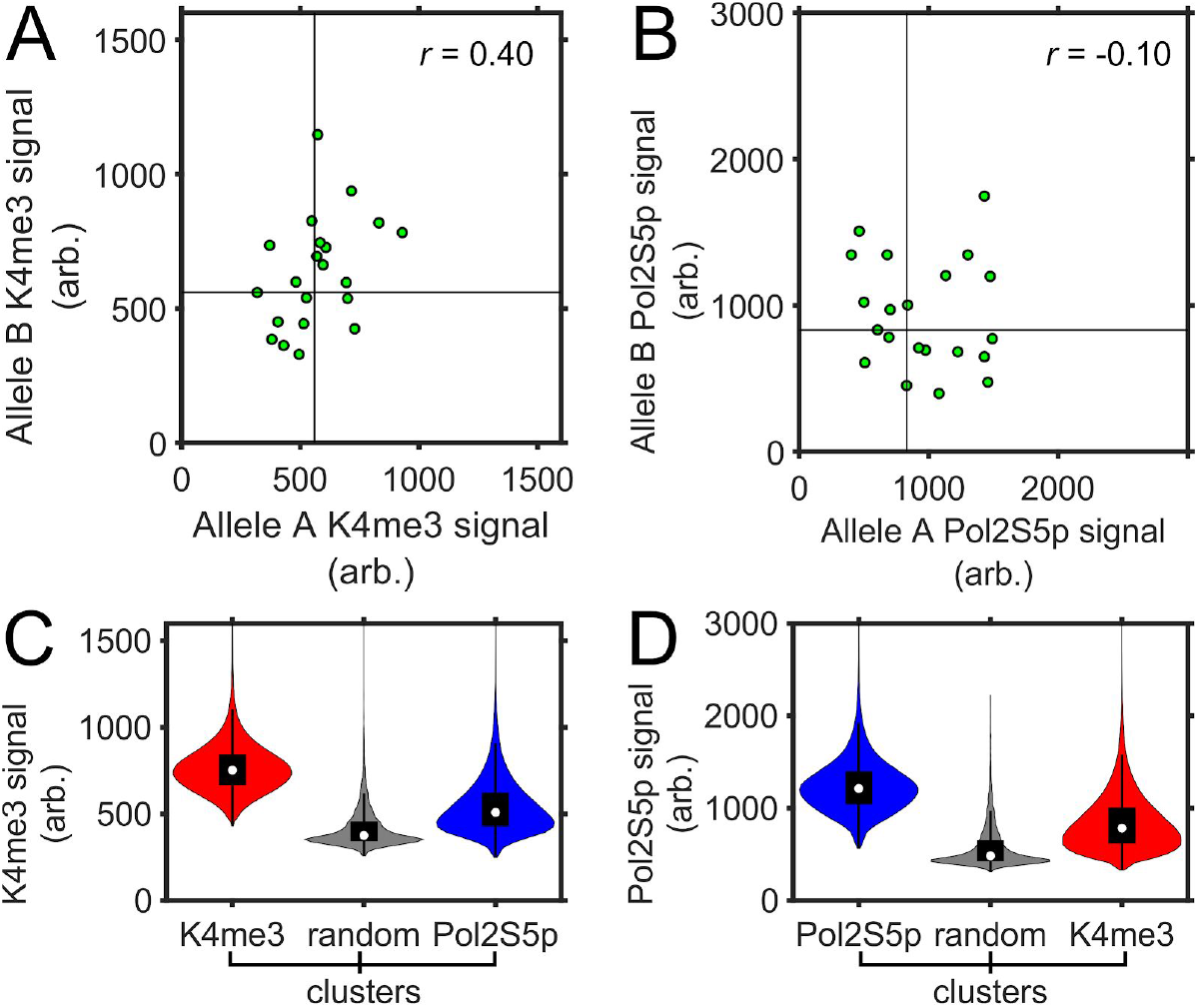
SCEPTRE compares H3K4me3 and paused RNA polymerase II signals between different *GAPDH* alleles in the same cell, or between immunolabeled cluster distributions. **(A-B)** Fluorescence signal (arb. = arbitrary units) of either H3K4me3 (K4me3) in **A.**, or paused RNA polymerase II (Pol2S5p) in **B.**, in *GAPDH* alleles within the same cell from the data set in **figure 5** (one locus from each cell containing 2-4 loci is randomly assigned as allele A, and a second locus as allele B). Black lines represent the threshold “on” level for each fluorescence signal. The correlation coefficient (r) is shown on the top-right corner of each plot. **(C-D)** Fluorescence signal (arb. = arbitrary units) for either H3K4me3 in **C.**, or paused RNA polymerase II in **D**., for each distribution of H3K4me3 (red), paused RNA polymerase II (blue) and randomly selected regions (random, gray) clusters within the cells in **figure 5**.

**Supplementary figure 11.**
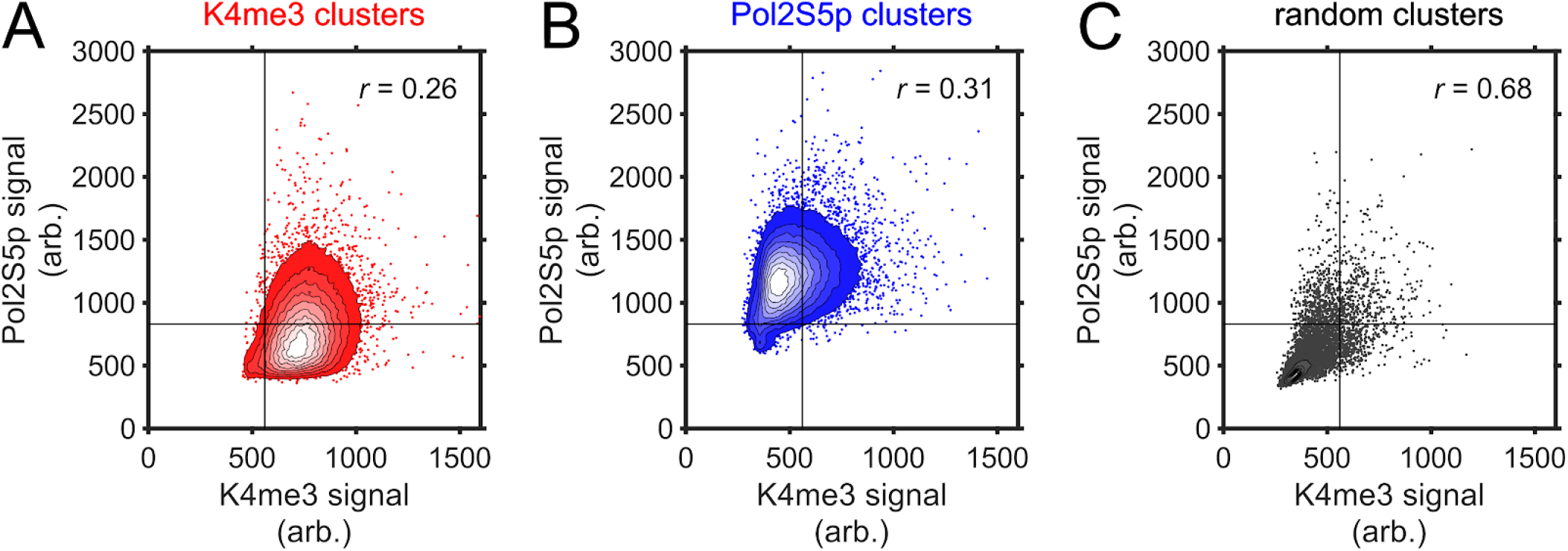
SCEPTRE compares H3K4me3 and paused RNA polymerase II signals within segmented immunostained and random clusters. Contours for the fluorescence signal (arb.=arbitrary units) frequency of H3K4me3 (K4me3) and paused RNA polymerase II (Pol2S5p) in the cluster sets of H3K4me3 (red) in **A.**, paused RNA polymerase II (blue) in **B.**, and randomly selected regions (random, gray) in **C**. Straight black lines represent the threshold “on” level for each fluorescence signal. Contours have uniformly spaced steps ranging from 0.1 to 0.9 frequency and represent all clusters obtained for cells in **figure 5**. The remaining scatter in **A.** and **B.** is a 100-fold downsample of the original data by random selection for plot representation purposes. Correlation coefficients (r) for each data set, which are calculated before downsampling, are shown in the top-right corner of each plot.

**Supplementary figure 12.**
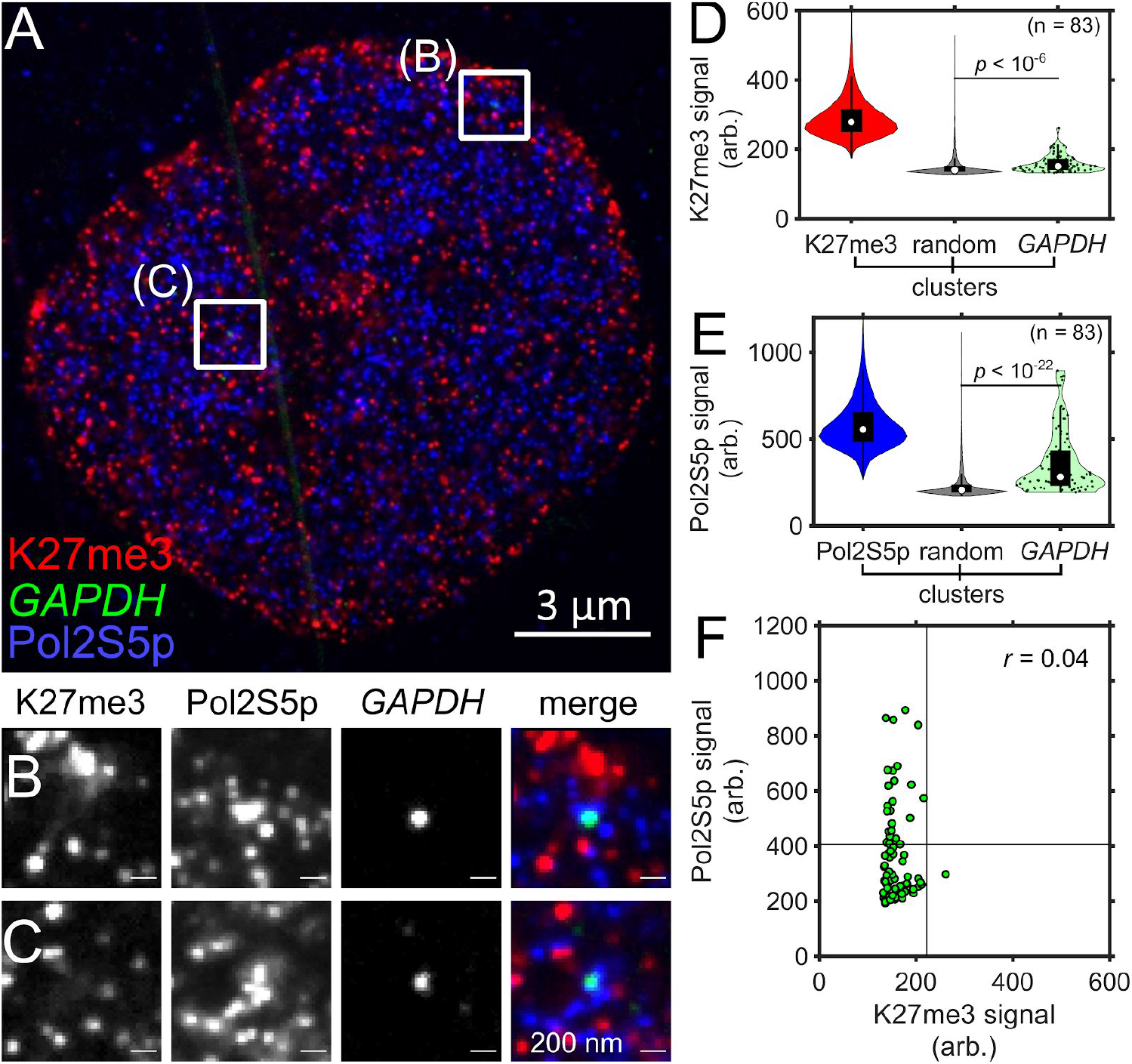
SCEPTRE distinguishes between H3K27me3 and paused RNA polymerase II signals at a single genomic region. **(A)** An expanded RPE1 cell with immunolabeled H3K27me3 (K27me3, red) and paused RNA polymerase II (Pol2S5p, blue), and FISH-labeled *GAPDH* (green). **(B,C)** Zoomed in views of the approximate center plane of an image stack for each *GAPDH* allele in the cell seen in **A. (D)** Distributions of H3K27me3 fluorescence signal (arb. = arbitrary units) within H3K27me3, randomly selected regions (random) and *GAPDH* clusters. **(E)** Distribution of paused RNA polymerase II fluorescence signal within paused RNA polymerase II, randomly selected regions and *GAPDH* clusters. **(F)** H3K27me3 and paused RNA polymerase II fluorescence signals within *GAPDH* clusters (green). Black lines represent the threshold “on” level for each fluorescence signal. Cluster numbers are K27me3 = 174072, Pol2S5p = 213724, random = 6099, *GAPDH* = 83. Significance determined by a right-tailed Wilcoxon rank-sum test of fluorescence signals in *GAPDH* against random cluster distributions. All scale bars are in pre-expansion units.

**Supplementary figure 13.**
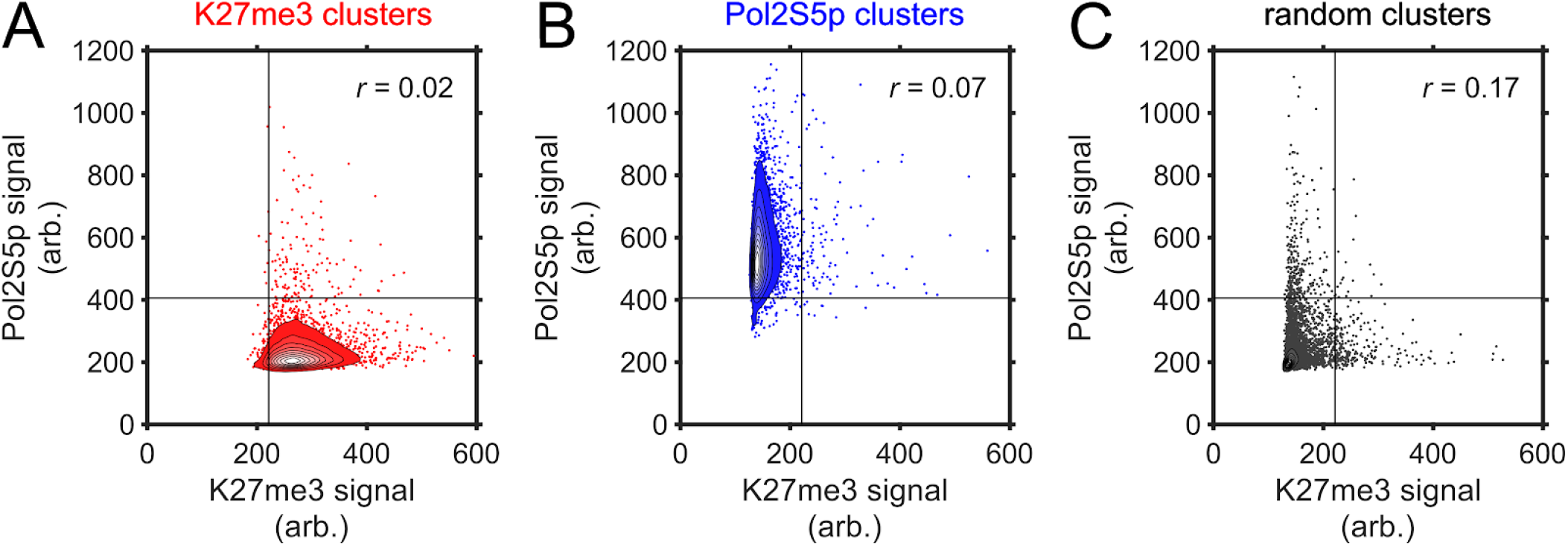
SCEPTRE compares H3K27me3 and paused RNA polymerase II signals within segmented immunostained and random clusters. Contours for the fluorescence signal (arb.=arbitrary units) frequency of H3K27me3 (K27me3) and paused RNA polymerase II (Pol2S5p) in the cluster sets of H3K27me3 (red) in **A.**, paused RNA polymerase II (blue) in **B.**, and randomly selected regions (random, gray) in **C**. Straight black lines represent the threshold “on” level for each fluorescence signal. Contours have uniformly spaced steps ranging from 0.1 to 0.9 frequency and represent all clusters obtained for cells in **supplementary figure 12**. The remaining scatter in **A.** and **B.** is a 100-fold downsample of the original data by random selection for plot representation purposes. Correlation coefficients (r) for each data set, which are calculated before downsampling, are shown in the top-right corner of each plot.

**Supplementary figure 14.**
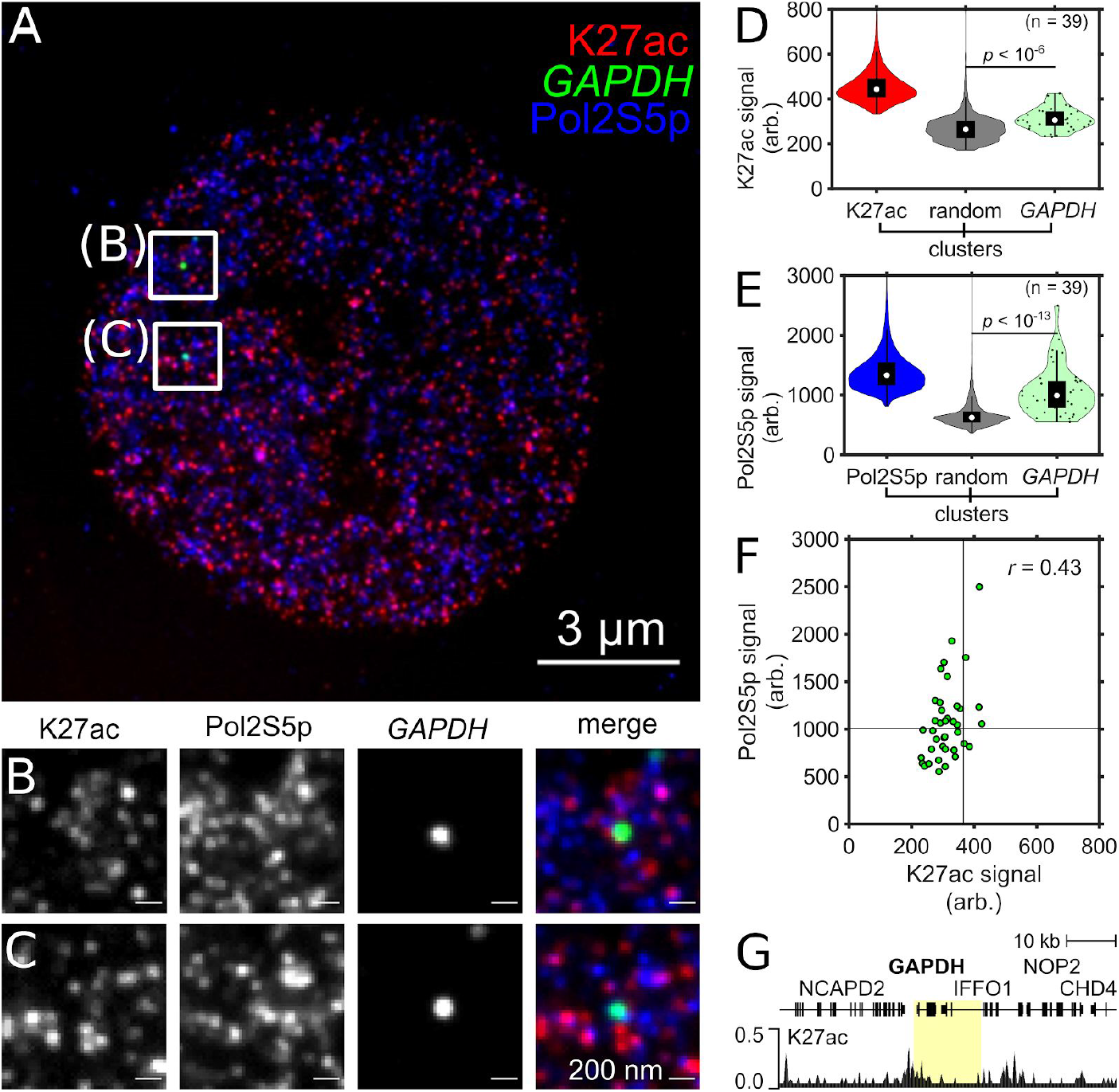
SCEPTRE compares H3K27ac and paused RNA polymerase II signals at a single genomic region. **(A)** An expanded RPE1 cell with immunolabeled H3K27ac (K27ac, red) and paused RNA polymerase II (Pol2S5p, blue), and FISH-labeled *GAPDH* (green). **(B,C)** Zoomed in views of the approximate center plane of an image stack for each *GAPDH* allele in the cell seen in **A. (D)** Distributions of H3K27ac fluorescence signal (arb. = arbitrary units) within H3K27ac, randomly selected regions (random) and *GAPDH* clusters. **(E)** Distribution of paused RNA polymerase II fluorescence signal within paused RNA polymerase II, randomly selected regions and *GAPDH* clusters. **(F)** H3K27ac and paused RNA polymerase II fluorescence signals within *GAPDH* clusters (green). Black lines represent the threshold “on” level for each fluorescence signal. (**G**) CUT&RUN normalized counts for H3K27ac marks in RPE1 cells for the FISH targeted *GAPDH* region (highlighted). Cluster numbers for **D.** and **E.** are K27ac = 82644, Pol2S5p = 153482, random = 3815, *GAPDH* = 39. Significance determined by a right-tailed Wilcoxon rank-sum test of fluorescence signals in *GAPDH* against random cluster distributions. All scale bars are in pre-expansion units.

**Supplementary figure 15.**
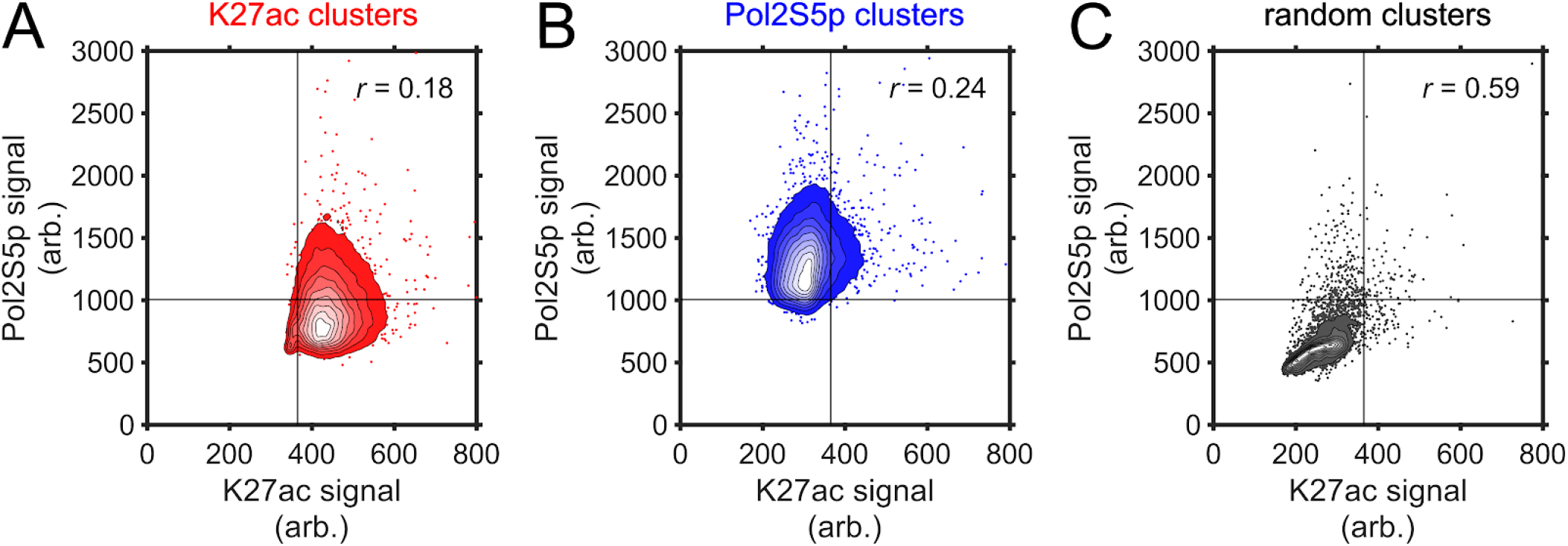
SCEPTRE compares H3K27ac and paused RNA polymerase II signals within segmented immunostained and random clusters. Contours for the fluorescence signal (arb.=arbitrary units) frequency of H3K27ac (K27ac) and paused RNA polymerase II (Pol2S5p) in the cluster sets of H3K27ac (red) in **A.**, paused RNA polymerase II (blue) in **B.**, and randomly selected regions (random, gray) in **C**. Straight black lines represent the threshold “on” level for each fluorescence signal. Contours have uniformly spaced steps ranging from 0.1 to 0.9 frequency and represent all clusters obtained for cells in **supplementary figure 14**. The remaining scatter in **A.** and **B.** is a 100-fold downsample of the original data by random selection for plot representation purposes. Correlation coefficients (r) for each data set, which are calculated before downsampling, are shown in the top-right corner of each plot.

## Supplementary tables

**Supplementary table S1.**
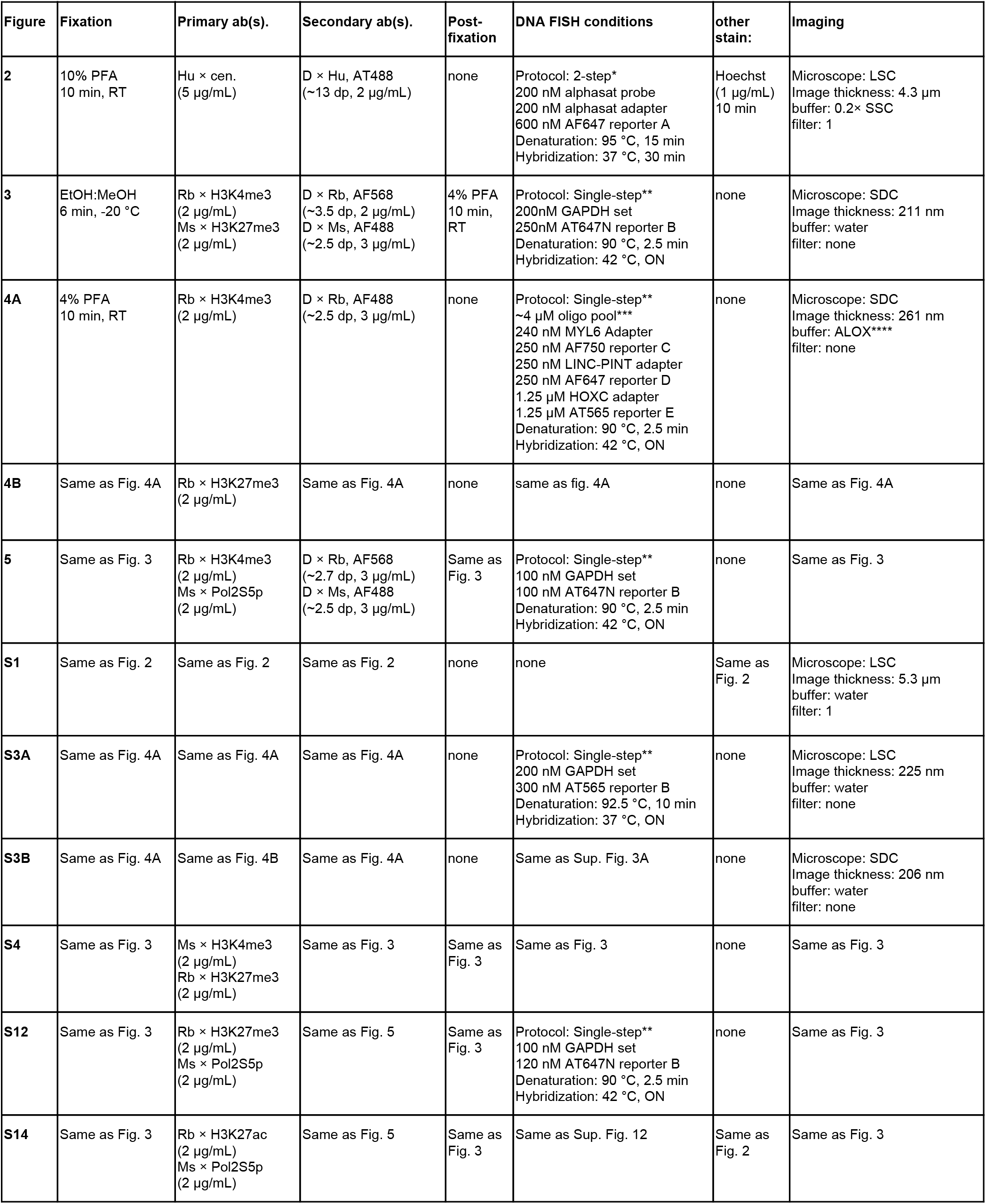
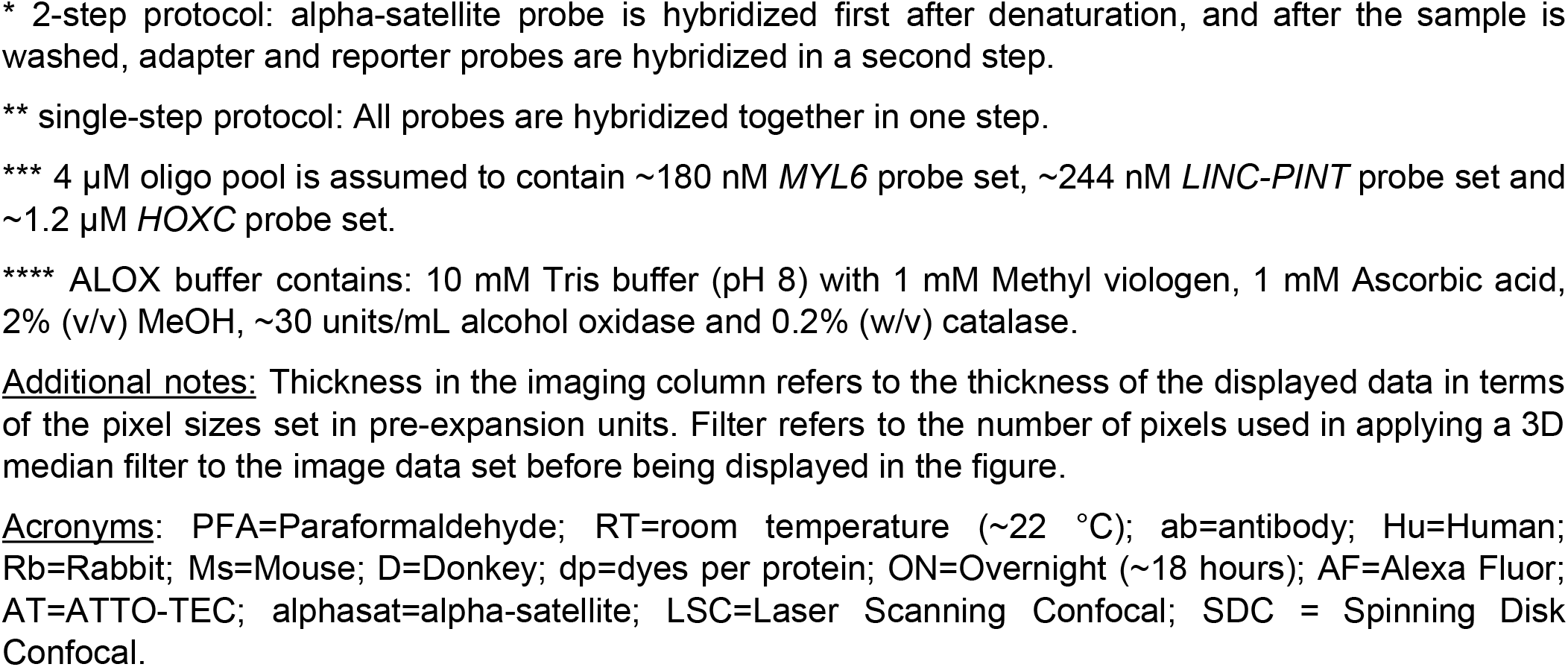
Summary of samp e preparation and imaging conditions.

| Figure | Fixation | Primary ab(s). | Secondary ab(s). | Post-fixation | DNA FISH conditions | other stain: | Imaging |
| --- | --- | --- | --- | --- | --- | --- | --- |
| 2 | 10% PFA<br>10 min, RT | Hu × cen.<br>(5 µg/mL) | D × Hu, AT488<br>(~13 dp, 2 µg/mL) | none | Protocol: 2-step*<br>200 nM alphasat probe<br>200 nM alphasat adapter<br>600 nM AF647 reporter A<br>Denaturation: 95 °C, 15 min<br>Hybridization: 37 °C, 30 min | Hoechst<br>(1 µg/mL)<br>10 min | Microscope: LSC<br>Image thickness: 4.3 µm<br>buffer: 0.2× SSC<br>filter: 1 |
| 3 | EtOH:MeOH<br>6 min, -20 °C | Rb × H3K4me3<br>(2 µg/mL)<br>Ms × H3K27me3<br>(2 µg/mL) | D × Rb, AF568<br>(~3.5 dp, 2 µg/mL)<br>D × Ms, AF488<br>(~2.5 dp, 3 µg/mL) | 4% PFA<br>10 min,<br>RT | Protocol: Single-step**<br>200nM GAPDH set<br>250nM AT647N reporter B<br>Denaturation: 90 °C, 2.5 min<br>Hybridization: 42 °C, ON | none | Microscope: SDC<br>Image thickness: 211 nm<br>buffer: water<br>filter: none |
| 4A | 4% PFA<br>10 min, RT | Rb × H3K4me3<br>(2 µg/mL) | D × Rb, AF488<br>(~2.5 dp, 3 µg/mL) | none | Protocol: Single-step**<br>~4 µM oligo pool***<br>240 nM MYL6 Adapter<br>250 nM AF750 reporter C<br>250 nM LINC-PINT adapter<br>250 nM AF647 reporter D<br>1.25 µM HOXC adapter<br>1.25 µM AT565 reporter E<br>Denaturation: 90 °C, 2.5 min<br>Hybridization: 42 °C, ON | none | Microscope: SDC<br>Image thickness: 261 nm<br>buffer: ALOX****<br>filter: none |
| 4B | Same as Fig. 4A | Rb × H3K27me3<br>(2 µg/mL) | Same as Fig. 4A | none | same as fig. 4A | none | Same as Fig. 4A |
| 5 | Same as Fig. 3 | Rb × H3K4me3<br>(2 µg/mL)<br>Ms × Pol2S5p<br>(2 µg/mL) | D × Rb, AF568<br>(~2.7 dp, 3 µg/mL)<br>D × Ms, AF488<br>(~2.5 dp, 3 µg/mL) | Same as<br>Fig. 3 | Protocol: Single-step**<br>100 nM GAPDH set<br>100 nM AT647N reporter B<br>Denaturation: 90 °C, 2.5 min<br>Hybridization: 42 °C, ON | none | Same as Fig. 3 |
| S1 | Same as Fig. 2 | Same as Fig. 2 | Same as Fig. 2 | none | none | Same as<br>Fig. 2 | Microscope: LSC<br>Image thickness: 5.3 µm<br>buffer: water<br>filter: 1 |
| S3A | Same as Fig. 4A | Same as Fig. 4A | Same as Fig. 4A | none | Protocol: Single-step**<br>200 nM GAPDH set<br>300 nM AT565 reporter B<br>Denaturation: 92.5 °C, 10 min<br>Hybridization: 37 °C, ON | none | Microscope: LSC<br>Image thickness: 225 nm<br>buffer: water<br>filter: none |
| S3B | Same as Fig. 4A | Same as Fig. 4B | Same as Fig. 4A | none | Same as Sup. Fig. 3A | none | Microscope: SDC<br>Image thickness: 206 nm<br>buffer: water<br>filter: none |
| S4 | Same as Fig. 3 | Ms × H3K4me3<br>(2 µg/mL)<br>Rb × H3K27me3<br>(2 µg/mL) | Same as Fig. 3 | Same as<br>Fig. 3 | Same as Fig. 3 | none | Same as Fig. 3 |
| S12 | Same as Fig. 3 | Rb × H3K27me3<br>(2 µg/mL)<br>Ms × Pol2S5p<br>(2 µg/mL) | Same as Fig. 5 | Same as<br>Fig. 3 | Protocol: Single-step**<br>100 nM GAPDH set<br>120 nM AT647N reporter B<br>Denaturation: 90 °C, 2.5 min<br>Hybridization: 42 °C, ON | none | Same as Fig. 3 |
| S14 | Same as Fig. 3 | Rb × H3K27ac<br>(2 µg/mL)<br>Ms × Pol2S5p<br>(2 µg/mL) | Same as Fig. 5 | Same as<br>Fig. 3 | Same as Sup. Fig. 12 | Same as<br>Fig. 2 | Same as Fig. 3 |

\* 2-step protocol: alpha-satellite probe is hybridized first after denaturation, and after the sample is washed, adapter and reporter probes are hybridized in a second step.
\*\* single-step protocol: All probes are hybridized together in one step.
\*\*\* 4 μM oligo pool is assumed to contain ~180 nM *MYL6* probe set, ~244 nM *LINC-PINT* probe set and ~1.2 μM *HOXC* probe set.
\*\*\*\* ALOX buffer contains: 10 mM Tris buffer (pH 8) with 1 mM Methyl viologen, 1 mM Ascorbic acid, 2% (v/v) MeOH, ~30 units/mL alcohol oxidase and 0.2% (w/v) catalase.
Additional notes: Thickness in the imaging column refers to the thickness of the displayed data in terms of the pixel sizes set in pre-expansion units. Filter refers to the number of pixels used in applying a 3D median filter to the image data set before being displayed in the figure.
Acronyms: PFA=Paraformaldehyde; RT=room temperature (~22 °C); ab=antibody; Hu=Human; Rb=Rabbit; Ms=Mouse; D=Donkey; dp=dyes per protein; ON=Overnight (~18 hours); AF=Alexa Fluor; AT=ATTO-TEC; alphasat=alpha-satellite; LSC=Laser Scanning Confocal; SDC = Spinning Disk Confocal.

**Supplementary table S2.** Summary of image processing and analysis conditions.

| Figure | nuclear mask channel | gaussian smooth (SD) | contrast adjustment threshold | binarization method | FISH size filter (voxels) |
| --- | --- | --- | --- | --- | --- |
| **2** | Hoechst | 2 | nuclear: 1<br>α-centromere: 2<br>anti-centromere: 5 | Otsu | α-cen. size: 20 – 10000 |
| **3** | H3K27me3 | 1 | nuclear: 3<br>GAPDH: 10<br>H3K4me3: 3<br>H3K27me3: 2 | Laplace | GAPDH: $\geq 50$ |
| **4A** | H3K4me3 | 1 | nuclear: 2.5<br>MYL6: 10<br>HOXC: 10<br>LINC-PINT: 10<br>H3K4me3: 2.5 | Laplace | MYL6: $\geq 20$<br>HOXC: $\geq 20$<br>LINC-PINT: $\geq 20$ |
| **4B** | H3K27me3 | 1 | nuclear: 3<br>MYL6: 10<br>HOXC: 10<br>LINC-PINT: 10<br>H3K4me3: 3 | Laplace | MYL6: $\geq 20$<br>HOXC: $\geq 50$<br>LINC-PINT: $\geq 50$ |
| **5** | Pol2S5p | 1 | nuclear: 3<br>GAPDH: 15<br>H3K4me3: 5<br>Pol2S5p: 5 | Laplace | GAPDH: $\geq 50$ |
| **S3A** | H3K4me3 | 1 | nuclear: 4<br>GAPDH: 15<br>H3K4me3: 3 | Laplace | GAPDH: $\geq 80$ |
| **S3B** | H3K27me3 | 1 | nuclear: 3:<br>GAPDH: 15<br>H3K27me3: 3 | Laplace | GAPDH: $\geq 80$ |
| **S4** | H3K27me3 | 1 | Nuclear: 3<br>GAPDH: 10<br>H3K4me3: 3<br>H3K27me3: 3 | Laplace | GAPDH: $\geq 50$ |
| **S12** | H3K27me3 | 1 | Nuclear: 3<br>GAPDH: 15<br>H3K27me3: 9<br>Pol2S5p: 9 | Laplace | GAPDH: $\geq 75$ |
| **S14** | Hoechst | 1 | Nuclear: 1<br>GAPDH: 10<br>H3K27ac: 2<br>Pol2S5p: 4 | Laplace | GAPDH: $\geq 20$ |

Acronym: SD=standard deviation
Additional notes: Contrast adjustment threshold represents the number of third quartiles above the median of an image stack histogram used to establish the threshold for clipping during contrast adjustment (see Materials and Methods for more details).

